# RNA mis-splicing in children with myotonic dystrophy is associated with physical function

**DOI:** 10.1101/2024.07.03.600889

**Authors:** Julia M. Hartman, Kobe Ikegami, Marina Provenzano, Kameron Bates, Amanda Butler, Aileen S. Jones, Kiera N. Berggren, Jeanne Dekdebrun, Marnee J. McKay, Jennifer N. Baldwin, Kayla M.D. Cornett, Joshua Burns, Michael Kiefer, Nicholas E. Johnson, Melissa A. Hale, the DMCRN

## Abstract

**Objectives:** Dysregulated RNA alternative splicing is the hallmark of myotonic dystrophy type 1 (DM1). However, the association between RNA mis-splicing and physical function in children with the most severe form of disease, congenital myotonic dystrophy (CDM), is unknown.

**Methods:** 82 participants (42 DM1 adults & 40 CDM children) with muscle biopsies and measures of myotonia, motor function, and strength were combined from five observational studies. Data were normalized and correlated with an aggregate measure of alternative splicing dysregulation, [MBNL]_inferred_ in skeletal muscle biopsies. Multiple linear regression analysis was performed to predict [MBNL]_inferred_ using clinical outcome measures alone. Similar analyses were performed to predict 12-month physical function using baseline metrics.

**Results:** Myotonia (measured via vHOT) was significantly correlated with RNA mis-splicing in our cross-sectional population of all DM1 individuals; CDM participants alone displayed no myotonia despite a similar range of RNA mis-splicing. Measures of motor performance and muscle strength were significantly associated with [MBNL]_inferred_ in our cohort of all DM1 individuals and when assessing CDM children independently. Multiple linear regression analyses yielded two models capable of predicting [MBNL]_inferred_ from select clinical outcome assessments alone in all subjects (adjusted R^2^ = 0.6723) or exclusively in CDM children (adjusted R^2^ = 0.5875).

**Interpretation:** Our findings establish significant correlations between skeletal muscle performance and a composite measure of alternative splicing dysregulation, [MBNL]_inferred,_ in DM1. The strength of these correlations and the development of the predictive models will assist in designing efficacious clinical trials for individuals with DM1, particularly CDM.

## Introduction

Myotonic dystrophy type 1 (DM1) is the most common form of muscular dystrophy with a prevalence of 1 in 2100^1^. DM1 is inherited in an autosomal dominant manner and is caused by a CTG trinucleotide repeat expansion (CTG_n_) in the 3’ untranslated region of the dystrophia myotonica protein kinase (*DMPK*) gene^2–4^. Although nearly every organ system can be affected, the core clinical features of DM1 include progressive distal muscle weakness, early onset cataracts, and myotonia (i.e., delayed muscle relaxation following contraction)^5,6^.

Congenital myotonic dystrophy (CDM) is the most severe form of DM1 and results from large, intergenerational (CTG)_n_ expansion between parent and child^7–9^. CDM presents at birth with symptoms of hypotonia, clubfoot, feeding difficulties, and respiratory distress^10–12^. In contrast to adults with DM1 who present with a consistent, progressive decline in muscle function over time, children with CDM exhibit a natural improvement and stabilization of motor function and muscle performance in early childhood. As children with CDM progress through adolescence and into early adulthood, symptoms become more consistent with adult-onset DM1^12,13^.

In DM1 the core pathogenesis is the sequestration of muscleblind-like (MBNL) RNA binding proteins by toxic, expanded CUG_n_ RNAs within nuclear aggregates termed foci. This leads to an overall reduction in the functional concentration of MBNL in affected tissues and subsequent dysregulation of RNA metabolism^14,15^. As critical regulators of fetal to adult mRNA isoform transitions, depletion of functional MBNL levels leads to global perturbations in splicing regulation and reversion of transcripts to the fetal isoform in affected tissues^16–20^. Numerous transcripts are mis-spliced, although only a select few have been linked to disease phenotypes in DM1 cell and animal models, including *CLCN1* and myotonia^21,22^, *SCN5A* and cardiac arrhythmia^23^, and *BIN1* and muscle weakness^24^.

While RNA mis-splicing of individual events has been correlated with ankle dorsiflexion performance and manual muscle testing in individuals with adult-onset DM1^25,26^, this relationship has not been replicated using other measures of physical performance. Additionally, no such associations have been evaluated in children with CDM. Previous work by our group has characterized global splicing dysregulation in skeletal muscle of DM1 adults and a cohort of children with CDM using a composite measure of MBNL-dependent splicing, [MBNL]_inferred_. This representative metric of free intracellular MBNL concentration strongly correlates with global splicing dysregulation as captured by total RNA sequencing ^27,28^.

Using this data, we aimed to investigate if transcriptome-wide RNA mis-splicing as measured by [MBNL]_inferred_ correlates with a wider breadth of functional measures assessing myotonia, motor function, and strength in a cross-sectional cohort of adults with DM1 and children with CDM. Additionally, we sought to evaluate if [MBNL]_inferred_ levels in individuals with CDM correlate with the patterns of functional improvement observed throughout childhood in this affected population. Using these correlative analyses, we developed regression models to predict [MBNL]_inferred_ using measures of physical function. This model may offer a non-invasive alternative for predicting disease associated mis-splicing in affected individuals. Lastly, we developed regression models to predict 12-month physical function using baseline performance and baseline [MBNL]_inferred_ values.

## Methods

### Study Design

Clinical phenotypes and associated biopsies from 5 multisite longitudinal observational studies were used to determine the correlation between splicing dysregulation (as measured by MBNL_inferred_) and physical function in a cohort of children with CDM and adults with DM1. Muscle biopsies were collected at baseline. Clinical assessments were completed at baseline, and/or at a 3-month visit, and/or at a 12-month visit depending on the study they were enrolled in. Participants provided informed consent before enrollment per the local site’s Institutional Review Board. For participants under 18, written informed consent was obtained from one parent and verbal assent from children over the age of 8.

Children with CDM between the ages of 0-17 (inclusive) were enrolled in the HELP-CDM, TREAT-CDM, or ASPIRE studies (NCT03059264, NCT05224778)^13^. Diagnostic criteria for CDM were defined as symptoms of myotonic dystrophy in the newborn period (<30 days) including hypotonia, respiratory distress, feeding difficulty, or clubfoot requiring hospitalization greater than 72 hours, and a genetic test confirming an expanded trinucleotide (CTG) repeat in the *DMPK* gene (CTG repeats greater than 200) or an affected mother. Exclusion criteria were described previously^13^.

Adults with DM1 over the age of 18 were enrolled in the HELP-DM1 or END-DM1 studies (NCT03981575). Participants were enrolled if they had a clinical diagnosis of DM1 or a positive genetic test confirming an expanded CTG repeat in the *DMPK* gene. They were excluded if they had symptomatic renal or liver disease, uncontrolled diabetes mellitus or thyroid disorders, or were pregnant. Mexiletine or other anti-myotonia agents were required to be stopped at least 72 hours prior to a study visit. Additional muscle biopsies that did not have associated clinical data were utilized from an in-house biorepository study to complement Figure 1.

**Figure 1:**
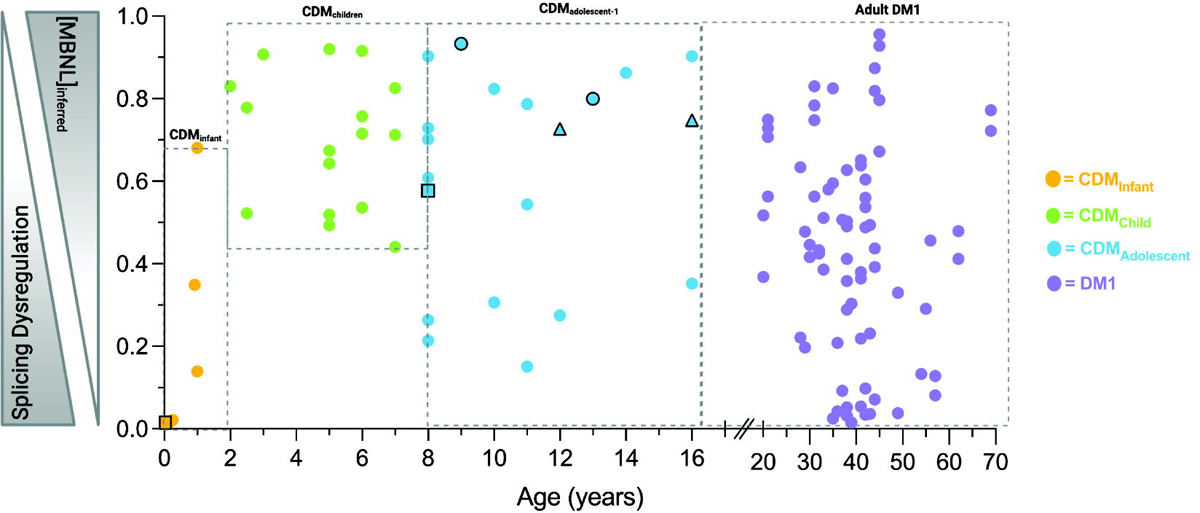
[MBNL]_inferred_ values calculated from DM1 and CDM skeletal muscle transcriptomes across pediatric development and adulthood. [MBNL]_inferred_ values range from 0-1 and inversely correlate with global mis-splicing severity. CDM infants (CDM_infant_; ≤ 2 yrs.) are shown in yellow ([MBNL]_inferred_ < 0.2). CDM children (CDM_child_; 2-8 yrs.) are shown in green ([MBNL]_inferred_ > 0.48), CDM adolescent (CDM_adolescent_; 8-16 yrs.) individuals are show in blue. Adult DM1 (20-69 yrs.) individuals are shown in purple. CDM-01 longitudinal biopsies shown as black-outlined squares. CDM-30 longitudinal biopsies shown as black-outlined circles. CDM-37 longitudinal biopsies are shown as black-outlined triangles.

### Clinical Assessments

Myotonia was assessed by participant (or caregiver if under 12) report and via clinical observation. Participants rated their myotonia severity on a 6-point scale from “1, I don’t experience this” to “6, It affects my life severely” as part of the Myotonic Dystrophy Health Index (MDHI) or Congenital and Childhood Myotonic Dystrophy Health Index (CCMDHI)^29^. Clinical myotonia was observed using video hand opening time (vHOT)^30,31^, in which the time to extend the thumb after 4 seconds of maximal hand flexion contraction is calculated via video recording of the procedure. The assessment was modified for children to squeeze a rubber toy for improved understanding of the task and time was scored live using a stopwatch.

Motor function measures included the 9-hole peg test^32^, 6-minute walk^33,34^, 4 stair climb^35^, and 10-meter walk/run test^35^ administered according to previously described methods. For adults enrolled in HELP-DM1, a 10-stair climb with railing as described in the modified dynamic gait index (mDGI) was performed^36^.

Strength measures for knee extension^37^, grip^13^, and ankle dorsiflexion^13^ were assessed via Quantitative Myometry (QMT) according to previously published methods. In adults, a fixed QMT system with a force transducer attached to a metal frame was used to provide adequate stabilization due to high force outputs. In children, handheld dynamometry was used to reduce the administration burden. Strength was measured in kilogram-force (kgf) units and converted to Newtons (N) by standard conversion of 9.80665 (e.g., 1 kgf = 9.80665 N).

### Muscle biopsy collection and derivation of [MBNL]_inferred_

Muscle biopsies from adult DM1 participants were collected with either a Bergstrom or 14-gauge argon Supercore needle of the tibialis anterior (TA) as previously described^38^. Muscle biopsies of the vastus lateralis from CDM children were obtained using either an open surgical technique or a 14-gauge Argon Supercore needle^28^. In all studies, biopsies were not performed in individuals with known bleeding disorders, a history of anti-coagulation medication use, or a platelet count less than 50,000. To participate in the muscle biopsy, ankle dorsiflexion strength had to be between 4+ and 4-on the Medical Research Council (MRC) scale for muscle strength.

Total RNA sequencing was performed on RNA extracted from all muscle biopsies and [MBNL]_inferred_ values were derived for each individual as previously described^27,28^. In brief, percent-spliced in (PSI) values from 9 skipped exon splicing events with high predictive power of overall global mis-splicing in DM1 muscle were used to calculate [MBNL]_inferred_^27,28^. [MBNL]_inferred_ values and CDM participants sub-cohort classifications were previously defined^28^. CDM sub-cohort classification of included individuals was defined based on age and [MBNL]_inferred_. The CDM sub-cohorts are defined as i) CDM_infant_ (≤ 2 years, [MBNL]_inferred_ < 0.7), ii) CDM_child_ (>2-8 years, [MBNL]_inferred_ > 0.4), iii) CDM_adolescent_ (≥ 8 years)^28^.

Individual PSI values for *CLCN1* exon 7a and *CACNA1S* exon 29 events were derived from comparisons between each affected group (CDM sub-cohort or DM1 adults) and associated age-matched, unaffected individuals as previously reported^28^.

### Statistical Analyses

Motor function and strength measures were converted to percent predicted of sex and age-matched healthy controls for analysis. Percent predicted for walk/running velocity was calculated via Pereira et al and Bohannon et al^39,40^ and percent predicted for the 6-minute walk test, stair climb velocity, 9-hole peg test, grip strength, ankle dorsiflexion, and knee extension were calculated using raw data from the 1000 Norms Project^41–43^.

Study data were collected and managed using REDCap® electronic data capture tools^44,45^ and exported to Microsoft Excel Version 16.75 for analysis. Statistical analyses were performed using GraphPad Prism 10.1.0 and p-values < 0.05 were considered significant. Clinical data points inconsistent with the cohort distribution were verified with the original data sources by the clinical evaluators and subsequently reviewed by the study’s Principal Investigator (P.I.).

Univariate correlations and multiple linear regression were performed in GraphPad Prism. Univariate Spearman correlations assumed data are not sampled from Gaussian distributions with a two tailed p-value and a 95% confidence interval. Given that myotonia was measured using a self-/parent-reporting scale, data was assumed to be non-parametric. Responses were accumulated between the various DM1 sub-groups and statistical significance between the group medians was calculated using a Kruskal-Wallis test. Dunn’s test was used to correct for multiple comparisons. To test for the overall difference between total adult DM1 and CDM myotonia scale responses, a Mann-Whitney test was used that assumed non-Gaussian distributed data with a two-tailed p-value.

Multiple linear regression analysis assumed a least squares regression and adjusted R-squared was used to quantify goodness-of-fit. D’Agostino-Pearson omnibus normality test, Anderson-Darling test, Shapiro-Wilk normality test, and the Kolmogorov-Smirnov normality test with Dallal-Wilkinson-Lillie for p-value were all performed alongside multiple linear regression analysis to ensure data normality. A 95% confidence level was used.

## Results

### Evaluation of [MBNL]_inferred_ in CDM children and DM1 adult skeletal muscle biopsies

We had previously used total RNA sequencing to characterize MBNL-dependent spliceopathy in skeletal muscle of CDM and DM1 subjects and calculated [MBNL]_inferred_, an aggregate metric of RNA mis-splicing representative of estimated intracellular concentrations of free MBNL. MBNL]_inferred_ values range from 0-1, with 0 indicating high MBNL depletion consistent with significant mis-splicing and 1 representative of intracellular MBNL concentrations comparable to unaffected individuals. Using this methodology we identified a triphasic modality of RNA mis-splicing progression across pediatric development within our cross-sectional cohort of CDM children which appeared to mirror the triphasic phenotypic progression observed^28^. In brief, CDM individuals under the age of 2 (CDM_infant_) presented with severe mis-splicing consistent with the severity of disease presentation at birth. This is followed by a universal and significant reversal of splicing dysregulation in early childhood (CDM_child_, 2 – 8 years). This observation is supported by the longitudinal sampling of CDM-01 at 2 weeks and 8 years of age where a marked increase in [MBNL]_inferred_ was observed ([MBNL]_inferred_ = 0.015 and 0.571, respectively). RNA mis-splicing universally improved in early childhood (CDM_child_, 2 – 8 years) with some individuals approaching splicing patterns like that of unaffected age-matched individuals. Sampled adolescent children displayed a spectrum of [MBNL]_inferred_ values, indicating a wide range of global splicing dysregulation. Post 8 years of age, a gradient of [MBNL]_inferred_ in CDM_adolescent_ was observed and mirrors that of adult DM1 individuals, for which a full range of splicing dysregulation is observed. This triphasic pattern of RNA mis-splicing throughout pediatric development in CDM children is visualized in Figure 1.

To assess potential correlations between skeletal muscle performance measures and alternative splicing dysregulation, we aggregated CDM and DM1 subjects that (i) received a muscle biopsy, (ii) had a [MBNL]_inferred_ score derived from matched skeletal muscle RNA, and (iii) possessed clinical outcomes from the same associated visit. While a total of 116 muscle biopsies were available from 82 unique participants across the five natural history studies and an in-house biorepository study (Figure 1, Supplemental Table 1), only 103 of the biopsies had associated clinical outcome data (n = 31 CDM & 72 DM1, Supplemental Table 2). The finalized cross-sectional cohort that we used for the analyses herein contains participants with time-point matched [MBNL]_inferred_ scores and clinical outcomes from either baseline visits (children and adults), 3-month visits (adults only), or sporadic longitudinal sampling (3 children with CDM only, indicated in Figure 1). Additional clinical information from participants at 12-month visits was collected for predictive modeling even though there was no associated muscle biopsy, and therefore no available [MBNL]_inferred_ score, from that timepoint (Supplemental Table 1 and 2). One sample (CDM-38) was excluded from subsequent data analysis as the biopsy was collected from the soleus. Baseline visit characteristics including mean [MBNL]_inferred_ and measures of myotonia, motor function, and strength are presented in Table 1 for all disease sub-cohorts. Raw clinical outcome data for all participants at all biopsy-matched timepoint sampling can be found in Supplemental Table 2.

**Table 1:** Characteristics of CDM and DM1 participants at baseline visit. Mean ± one SD reported unless otherwise indicated. n = number of samples, s = seconds, m = meters, kgf = kilogram-force.

| Baseline Characteristics | Infant | CDM<br>Child | Adolescent | DM1<br>Adults |
| --- | --- | --- | --- | --- |
| **Sample Size (n)** | 7 | 16 | 21 | 41 |
| **Biological Sex (n)** |  |  |  |  |
| Male | 6 | 12 | 9 | 17 |
| Female | 1 | 4 | 7 | 24 |
| **Age (yrs.)** | 0.7±0.69 | 5±1.6 | 11.1±3 | 39.5±10.3 |
| **CTG repeat length** | 1600±141 | 1035±335 | 1210±526 | 426±254 |
| **[MBNL]<sub>inferred</sub>** | 0.2±0.27 | 0.7±0.16 | 0.58±0.28 | 0.46±0.24 |
| **Biopsy Site** | Vastus lateralis | Vastus lateralis | Vastus lateralis | Tibialis anterior |
| **Clinical Outcome Measures** |  |  |  |  |
| **Myotonia Scale** | 1 | 2.1±1.4 | 2.4±1.5 | 3.2±1.5 |
| **Myotonia vHOT<sub>thumb</sub> (s)** | NA | 0.46±0.2 | 1.1±1.07 | 7.5±8.4 |
| **6-minute walk (m)** | NA | 338±91 | 3±137 | 377±103 |
| **Walk/running velocity (m/s)** | NA | 1.9±0.61 | 2.1±0.71 | 1.99±0.94 |
| **Stair climb velocity (stairs/s)** | NA | 1.16±0.43 | 1.26±0.70 | 1.7±0.68 |
| **9-Hole Peg Test (s)** | NA | 47.7±14.7 | 42.6±31 | 20±6.6 |
| **Grip strength (kgf)** | NA | 4.12±1.5 | 7.5±4.9 | 13.9±8.5 |
| **Ankle dorsiflexion (kgf)** | NA | 4.5±2.6 | 5.8±3.5 | 8.07±5.4 |
| **Knee Extension (kgf)** | NA | 8.2±2.6 | 9.8±3.8 | 22.1±9.0 |

### Alternative splicing metrics correlate with myotonia measures in all DM1 participants but not in CDM children alone

Myotonia was assessed via two methods - a qualitative participant/caregiver reported score of impact on daily living (from MDHI/CCHDMI) and quantitatively via video of hand opening time (vHOT). The impact of myotonia on a participant’s life via participant/caregiver reported scale was found to be significantly different between all groups (Figure 2a). Given the large observed differences in mean [MBNL]_inferred_ values between CDM sub-cohorts (Table 1), it is reasonable to infer that [MBNL]_inferred_ values correlate minimally with the impact of myotonia on a participant’s life (Figure 2a). Multiple comparisons testing revealed a significant difference between CDM_child_ responses (median response ≅ 1) and adult DM1 responses (median response ≅ 3) to the myotonia scale (p = 0.0498) (Figure 2a). When comparing the values of the self - reported scale in adult DM1 individuals to total CDM responses (CDM_total_), a significant increase in perceived impact of myotonia was observed for adults with DM1 (Figure 2b).

**Figure 2:**
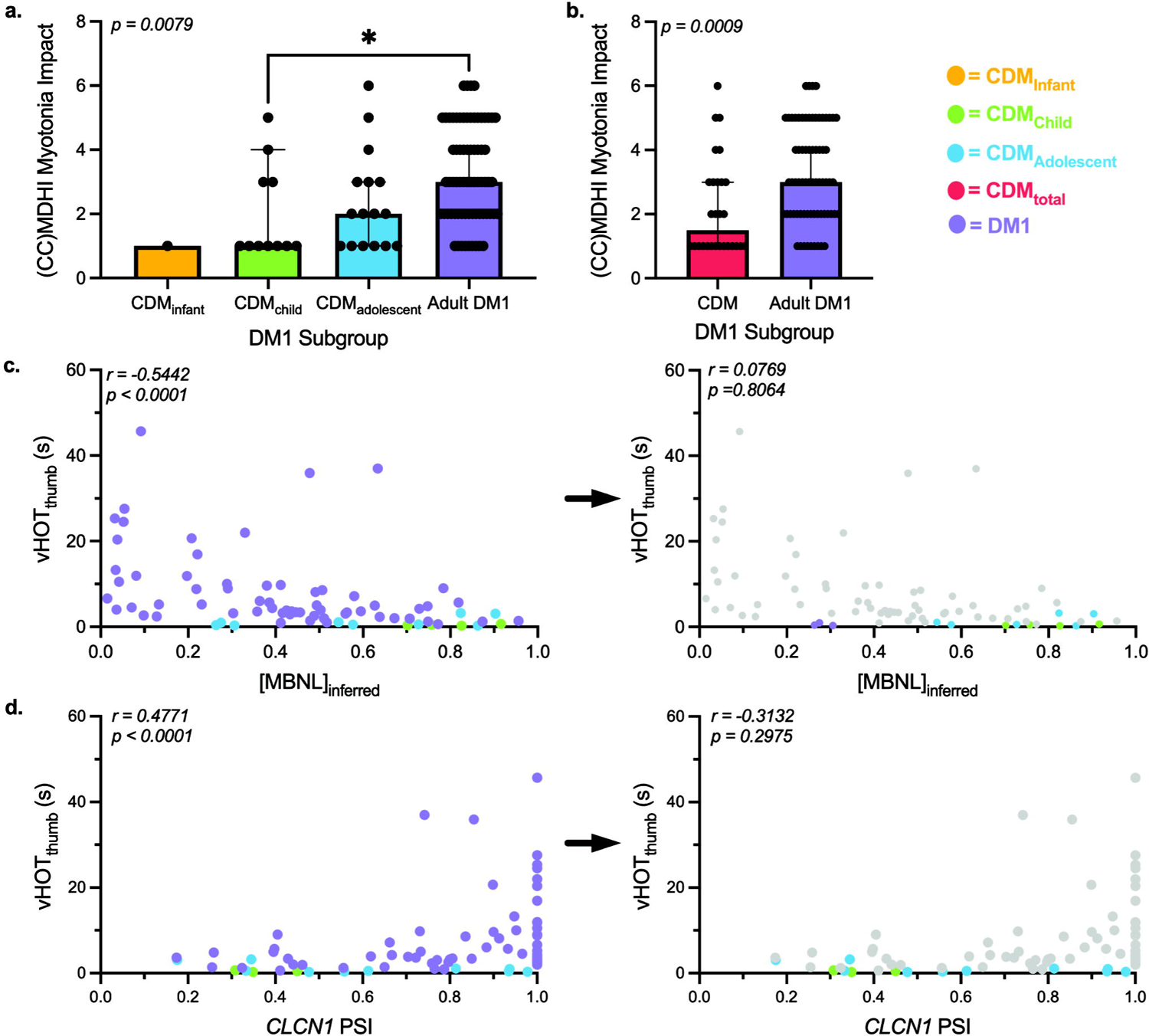
Myotonia measures in adult DM1 and CDM participants correlate disparately with skeletal muscle spliceopathy. (a.) Myotonia MDHI/CCMDHI scale average responses across CDM sub-cohorts and adult DM1 participants. Results are expressed as median ± 95% confidence interval (CI) via Kruskal-Wallis test with Dunn’s multiple comparisons test. (b.) Myotonia scale average value between DM1 individuals and all CDM individuals. Results are expressed as median ± 95% CI via unpaired Mann-Whitney test. (c.) Correlation between [MBNL]_inferred_ and vHOT_thumb_ times in all DM1 individuals (left) and CDM individuals alone (right panel). Adult DM1 measures are removed from the statistical analysis in the right panel but greyed out and included in the figure for visual reference. (d.) Correlation between *CLCN1* exon 7a percent spliced in (PSI) and vHOT_thumb_ times in all DM1 individuals and CDM individuals. All correlations are reported from a two-tailed Spearman test.

vHOT was used to quantitatively evaluate clinical myotonia between our DM1 subgroups. vHOT_thumb_ was found to correlate moderately with [MBNL]_inferred_ in all DM1 individuals (Spearman r = −0.54) (Figure 2c, left). However, there was no significant correlation observed in CDM participants alone despite the comparable range of spliceopathy observed (Spearman r = 0.08)(Figure 2c, right). To further investigate this phenomenon, we directly examined the relationship between vHOT and *CLCN1* exon 7a inclusion (percent spliced in, PSI). Defects in skeletal muscle chloride conductance due to mis-splicing of *CLCN1* are postulated to be causative of myotonia and restoration of aberrant exon 7a inclusion corrects this phenotype in DM1 mouse models^46^. Consistent with this pathogenic mechanism, *CLCN1* PSI was significantly correlated with vHOT_thumb_ times in all DM1 individuals. In contrast, this relationship was not observed in CDM participants alone (Figure 2d). While select CDM_adolescent_ individuals have nearly 100% *CLCN1* exon 7a inclusion, like that of their adult DM1 counterparts with the highest vHOT_thumb_ times (PSI > 0.8), minimal myotonia was observed. Mis-splicing of *CACNA1S* (CaV1.1), a calcium channel that controls skeletal muscle excitation–contraction coupling, has also been associated with exacerbated myopathy and myotonia in DM1^47,48^. In our complete cross-sectional cohort, reduced *CACNA1S* exon 29 PSI was significantly correlated with longer vHOT_thumb_ times, but not in CDM participants alone (Supplementary Figure 1). Overall, the lack of correlation between CDM vHOT_thumb_ times and either a composite measure of disease-associated spliceopathy, such as MBNL_inferred,_ or mis-splicing of specific, phenotype-associated events, such as CLCN1 exon 7a, is driven by the lack of quantifiable myotonia within children with CDM throughout development. This is consistent with previous observations that CDM individuals within the first decade of life do not experience clinical myotonia^49^. However, our analysis is the first of its kind to demonstrate that this phenomenon occurs even when phenotype-associated events remain mis-spliced at levels comparable to DM1 adults with severe myotonia.

### RNA mis-splicing correlates moderately with select motor function measures in all DM1 participants

We next assessed whether global mis-splicing as measured by [MBNL]_inferred_ correlated with several timed motor tests within all DM1 participants or selectively within CDM children, as utility of certain clinical outcome measures may vary with age and disease severity^50^.

The 9-hole peg test was significantly correlated, albeit weakly, with [MBNL]_inferred_ values in all DM1 individuals (Spearman r = −0.22). The relative association significantly improved in CDM children alone whereby individuals with higher [MBNL]_inferred_ were able to perform the test much quicker with results more in line with control times (i.e. resulting in a lower % predicted) (Spearman r = −0.66) (Figure 3a). In contrast to the 9-hole peg test, weak to moderate correlations were observed for both 6-minute walk distance and stair climbing velocity in all DM1 participants and in CDM children alone (Figure 3b and 3c). The strongest correlation observed between any motor function measure and [MBNL]_inferred_ was walk/running velocity Percent predicted walk/running velocity had a strong association with [MBNL]_inferred_ levels in all DM1 individuals that was only minimally reduced in CDM participants independently (Figure 3d).

**Figure 3:**
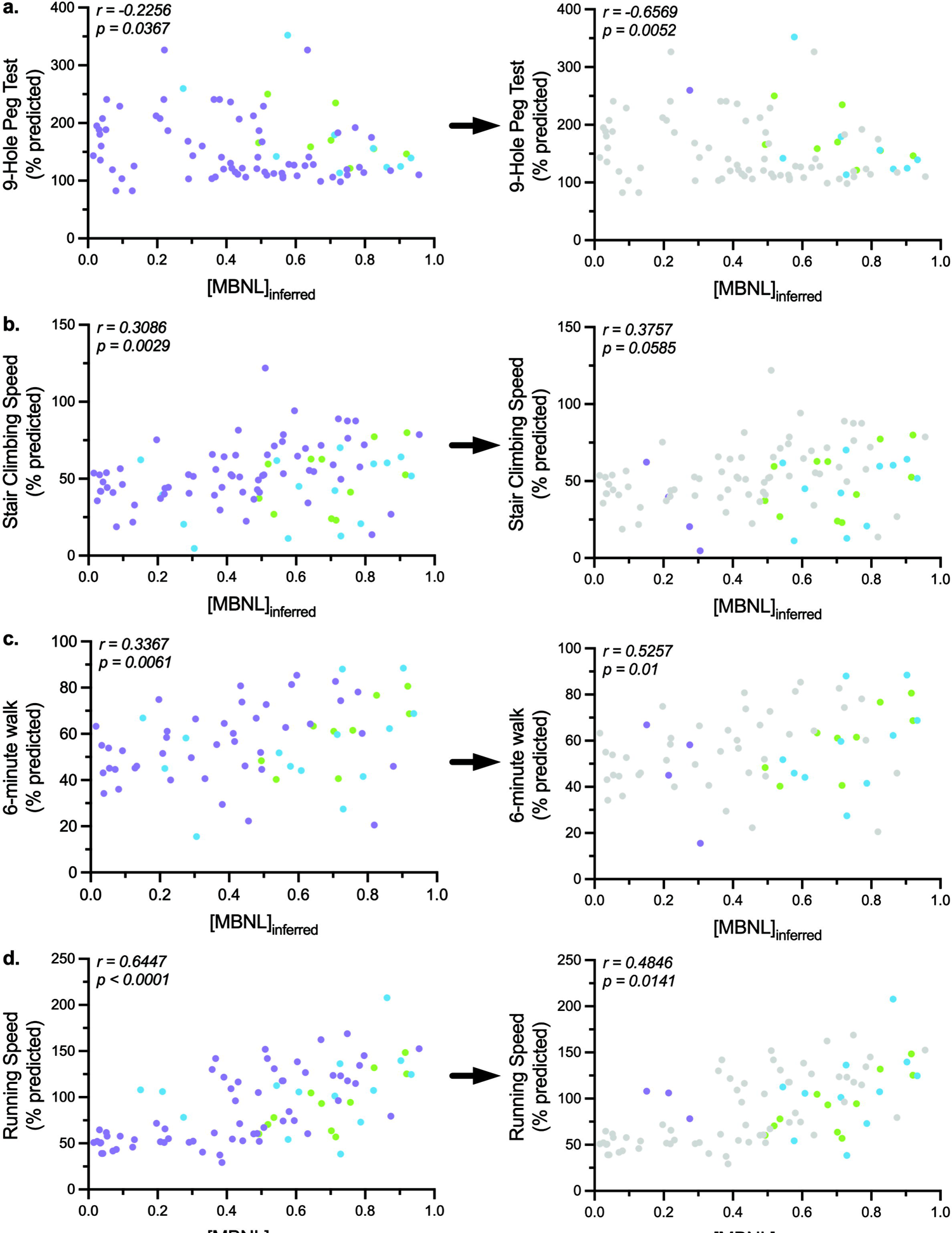
MBNL-dependent mis-splicing as measured by [MBNL]_inferred_ correlates moderately with motor function measures in DM1individuals. Correlation between [MBNL]_inferred_ and (a.) 9-hole peg test (% predicted), (b.) stair climbing velocity (% predicted), (c.) 6-minute walk distance (% predicted), and (d.) walk/running velocity (% predicted) in all DM1 individuals and CDM individuals. Left panels represent correlations of all DM1 subjects and right panels represent CDM subjects only. The adult DM1 measures were excluded from the CDM statistical analyses but greyed out and included in the figure on the right for visual reference. Correlations are reported from a two-tailed Spearman test.

### [MBNL]_inferred_ correlates strongly with skeletal muscle strength measures in all DM1 participants

Quantitative muscle testing of knee extension (KE), hand-grip (HGS), and ankle dorsiflexion (ADF) strength had the strongest associations with [MBNL]_inferred_ levels compared to all other outcome assessments utilized in these analyses. When DM1 adults and CDM children were evaluated in combination, higher levels of [MBNL]_inferred_ were strongly correlated with muscle strength; the highest reported association was with ADF (Spearman r = 0.68) (Figure 4). The strength of these relationships were generally maintained in CDM alone (Figure 4). Strikingly, the relative association of [MBNL]_inferred_ with ADF in CDM children was identical to that observed in all DM1 participants (Spearman r = 0.69), in part due to the range of ADF performance captured in CDM_adolescent_ participants that replicated the range observed in adults (Figure 4c, purple and blue samples). Consistent with correlative reports between RNA mis-splicing and muscle strength previously reported, measures of strength correlate strongly with [MBNL]_inferred_ in this assembled DM1 cohort.

**Figure 4:**
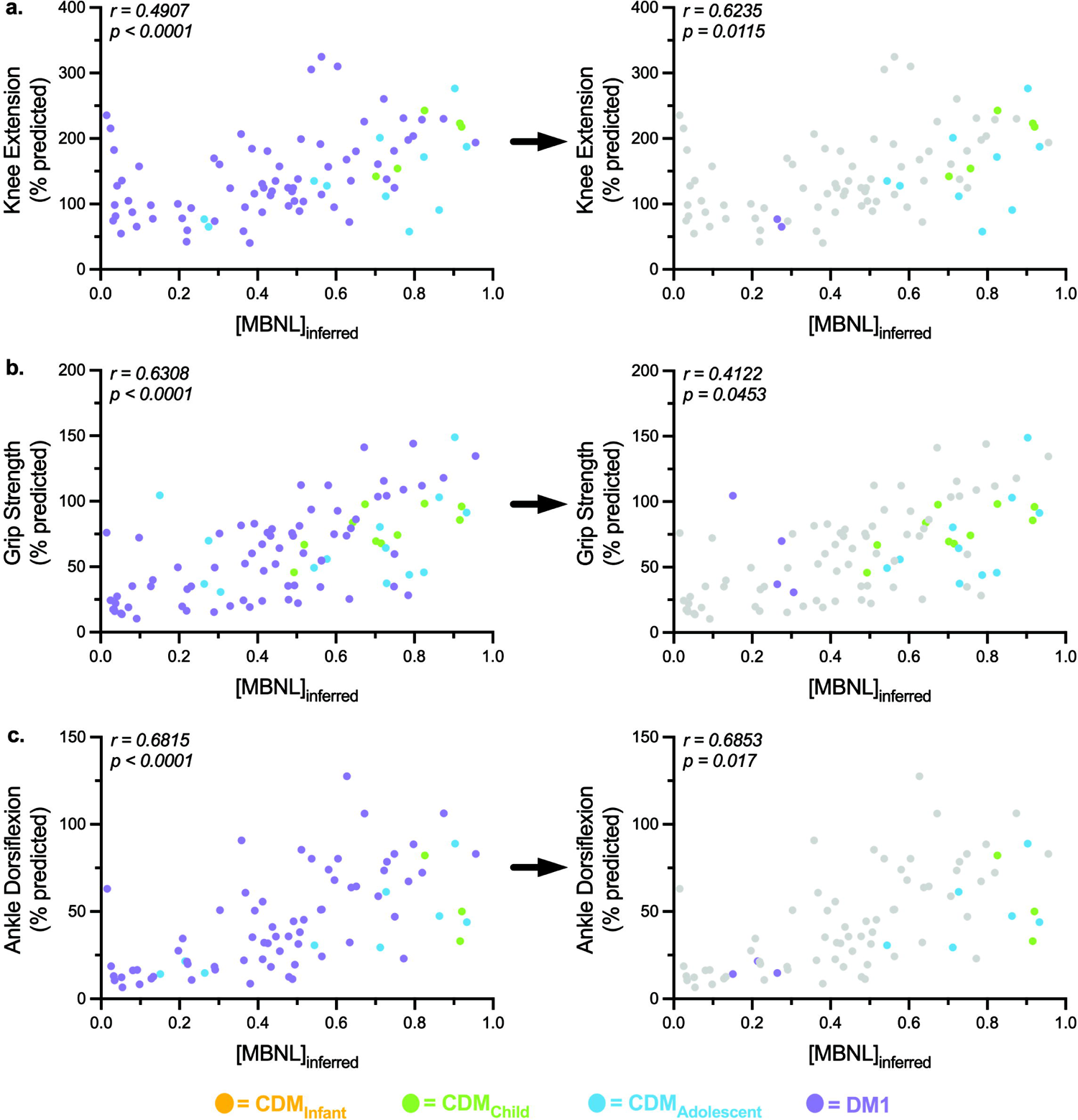
MBNL-dependent mis-splicing as measured by [MBNL]_inferred_ correlates with muscle strength independent of age. Correlation between [MBNL]_inferred_ and (a.) knee extension (% predicted), (b.) hand grip strength (% predicted), and (c.) ankle dorsiflexion (% predicted) in all DM1 individuals and CDM individuals. Left panels represent correlations of all DM1 subjects and right panels represent CDM subjects only. The adult DM1 measures were excluded from the CDM statistical analyses but greyed out and included in the figure on the right for visual reference. Correlations are reported from a two-tailed Spearman test.

### MBNL]_inferred_ can be predicted using physical function measures

Given the numerous significant correlations between [MBNL]_inferred_ and many of the clinical outcomes assessed above, we sought to determine if clinical outcome measures could accurately predict [MBNL]_inferred_ via multiple regression modeling. Measures significantly correlated with [MBNL]_inferred_ in the complete DM1 cohort inclusive of both adults and children (See Figures 2-4) were utilized in the “All – DM1 and CDM” model. Multiple linear regression analysis using all available sampled outcome measures led to a model that included 46 participants and had an adjusted r^2^ = 0.6723 whereby the developed model was able to account for majority of the variance in [MBNL]_inferred_ (Figure 5a, 5c).

**Figure 5:**
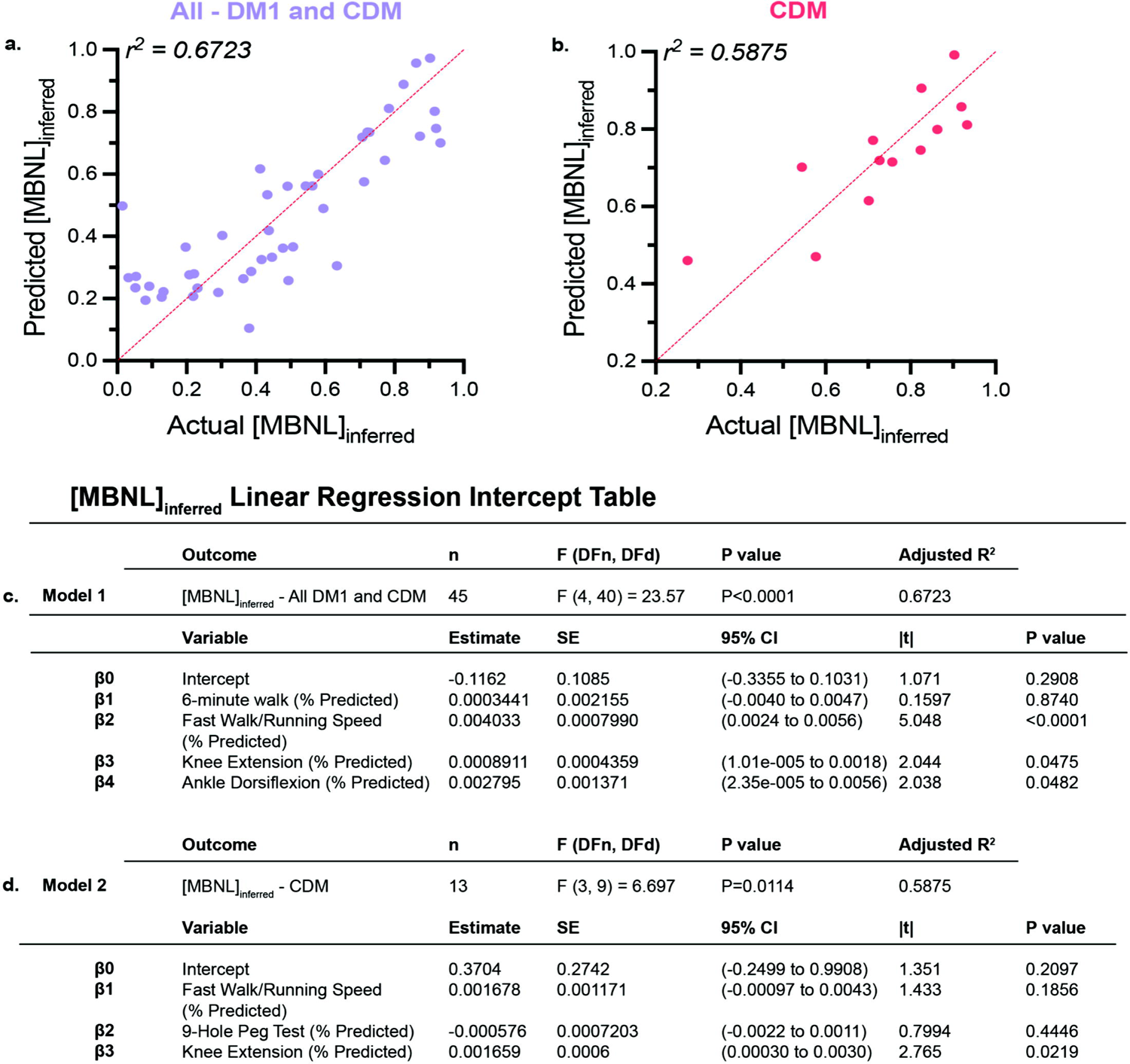
Multiple linear regression modeling to predict. [MBNL]_inferred_ *in DM1 participants using functional outcomes.* (a.) Multiple linear regression plot showing observed [MBNL]_inferred_ versus predicted [MBNL]_inferred_ for all DM1 individuals. (b.) Multiple linear regression plot showing observed [MBNL]_inferred_ versus predicted [MBNL]_inferred_ for CDM individuals. Intercept table of multiple linear regression models for (c.) complete DM1 cohort and (d.) CDM cohort alone using clinical outcome measures to predict [MBNL]_inferred_. Multiple statistical elements are reported (F = F-distribution (DFn = degrees of freedom numerator, DFd = degrees of freedom denominator), SE = standard error, 95% CI = 95% confidence interval, |t| = t-test statistic). Extended intercept tables for the complete DM1 cohort model are provided in Supplemental Table 3 and for the CDM cohort in Supplemental Table 4.

Next, we sought to build an equation to accurately predict [MBNL]_inferred_ in only children with CDM utilizing significantly correlated clinical outcome measures when this participant group was evaluated independently (See Figures 2-4). Multiple linear regression analysis using all available sampled CDM outcome measures led to a model that included 13 CDM participants and had an adjusted r^2^ = 0.5875 (Figure 5b-d). Despite the reduced sample size, the model still managed to moderately predict [MBNL]_inferred_ in children with CDM. Overall, these analyses indicate that clinical outcome measures of myotonia, motor function, and strength correlate significantly with MBNL dependent mis-splicing within all DM1 participants independent of age of disease onset. Additionally, these measures can be used in patients with DM1 to accurately predict disease-associated patterns of mis-splicing in skeletal muscle.

### 12-month physical function can be predicted using baseline physical function measure and baseline [MBNL]_inferred_

Given the strength of the correlations between [MBNL]_inferred_ and several of the clinical outcomes assessed above, we aimed to determine if future clinical performance could be accurately predicted using baseline [MBNL]_inferred_ values and baseline clinical performance in adults with DM1 and in children with CDM. We first looked at the outcome measures that had the strongest correlation with [MBNL]_inferred_ values within the complete cross-sectional cohort and CDM children alone – walk/running velocity and ADF strength.

Multiple linear regression analysis for predicting 12-month ambulation velocity using baseline walk/running velocity and [MBNL]_inferred_ in our cohort of adults with DM1 had an adjusted r^2^ = 0.681 (Figure 6a, Model 1). Modeling in CDM children alone suggested that these baseline values were similarly predictive of 12-month performance in CDM children alone (adjusted r^2^ = 0.646) (Figure 6a, Model 2). Interestingly, while baseline [MBNL]_inferred_ did not contribute significantly to the predictive power of either model, it was trending towards significance in the model for adults, but not for CDM subjects (p = 0.1675 & p = 0.7125, respectively) Multiple linear regression analysis for predicting 12-month ADF strength using baseline ankle dorsiflexion strength and [MBNL]_inferred_ in our cohort of adults with DM1 had an adjusted r^2^ = 0.7126 (Figure 6b, Model 1). The predictive power of the same model for CDM children was reduced (adjusted r^2^ = −0.4904) (Figure 6b, Model 2). Again, when we evaluated the contribution of baseline [MBNL]_inferred_ to our 12-month ADF predictive model in children with CDM, it was much less significant (p = 0.9233) as compared to that for adults with DM1 (p = 0.0895).

**Figure 6:**
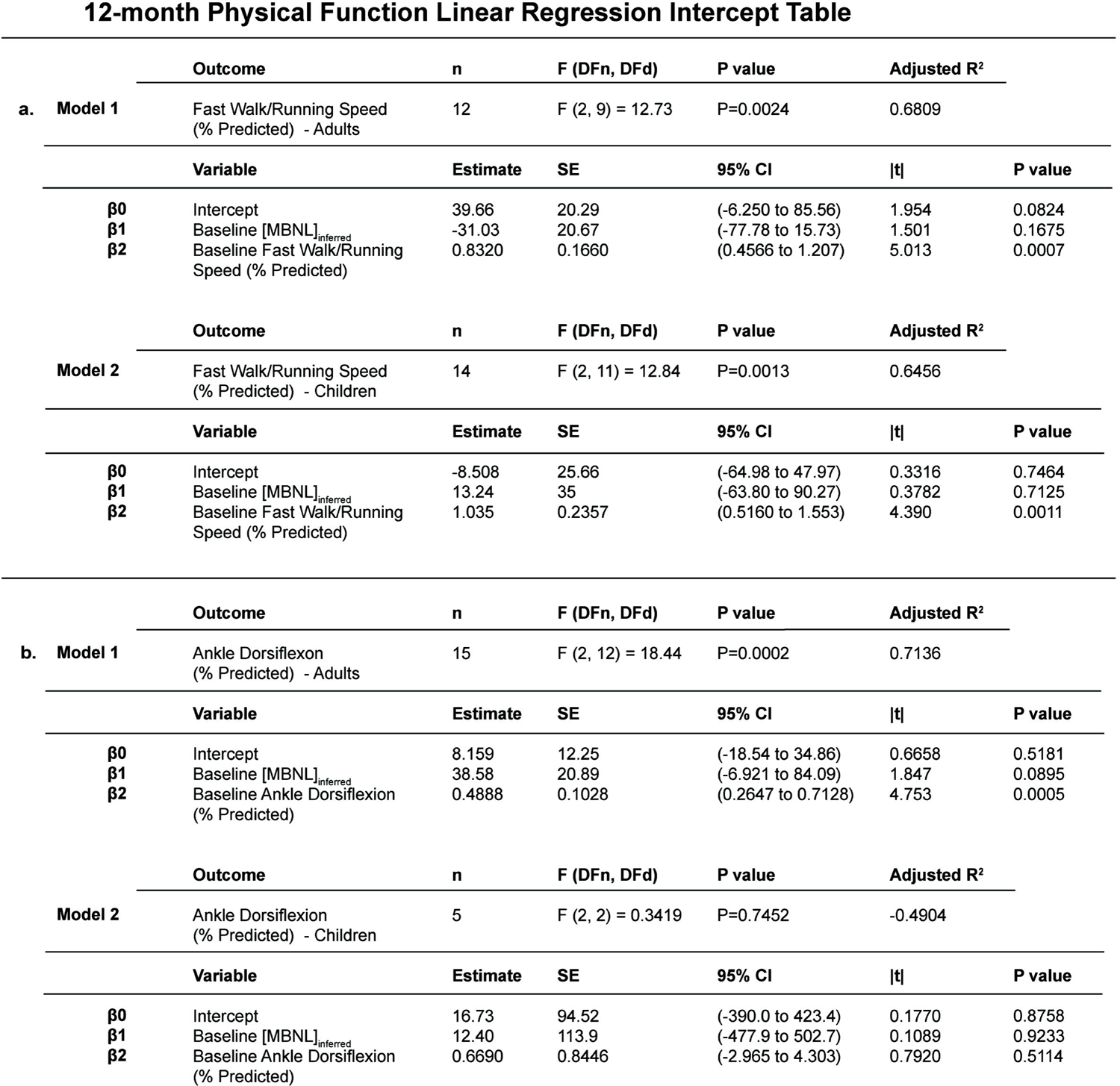
Multiple linear regression predictive models of 12-month ankle dorsiflexion strength and walk/running velocity in DM1 participants. Intercept table of multiple linear regression models for (a.) 12-month walk/running velocity (% predicted) and (b.) 12-month ankle dorsiflexion strength (% predicted) using baseline [MBNL]_inferred_ values and baseline physical function values. Multiple statistical elements are reported (F = F-distribution (DFn = degrees of freedom numerator, DFd = degrees of freedom denominator), SE = standard error, 95% CI = 95% confidence interval, |t| = t-test statistic). Extended intercept tables for the 12-month walk/running velocity models and 12-month ankle dorsiflexion strength are in Supplemental Table 5 and Supplemental Table 6, respectively.

Overall, of the seven motor function and strength outcome measures assessed, [MBNL]_inferred_ only contributed significantly to the predictive power of our 12-month multiple linear regression models for stair climbing velocity (Figure 7). In our complete cross-sectional cohort of both adults and children, baseline performance on this outcome assessment in combination with [MBNL]_inferred_ were able to accurately predict 12-month outcomes (adjusted r^2^ = 0.8057) (Figure 7a and 7b, Model 1). Multiple linear regression analysis for predicting 12-month stair climbing velocity using baseline stair climbing velocity values and baseline [MBNL]_inferred_ values in our cohort of CDM children alone had an adjusted r^2^ = 0.8629 (Figure 7b, Model 2), whereas the model for our cohort of adults with DM1 had an adjusted r^2^ = 0.6899 (figure 7b, Model 3). [MBNL]_inferred_ values predictive utility followed the same pattern observed above, whereby they were contributing more to our adult model (p = 0.0661) as compared to the model for children with CDM (p = 0.1049).

**Figure 7:**
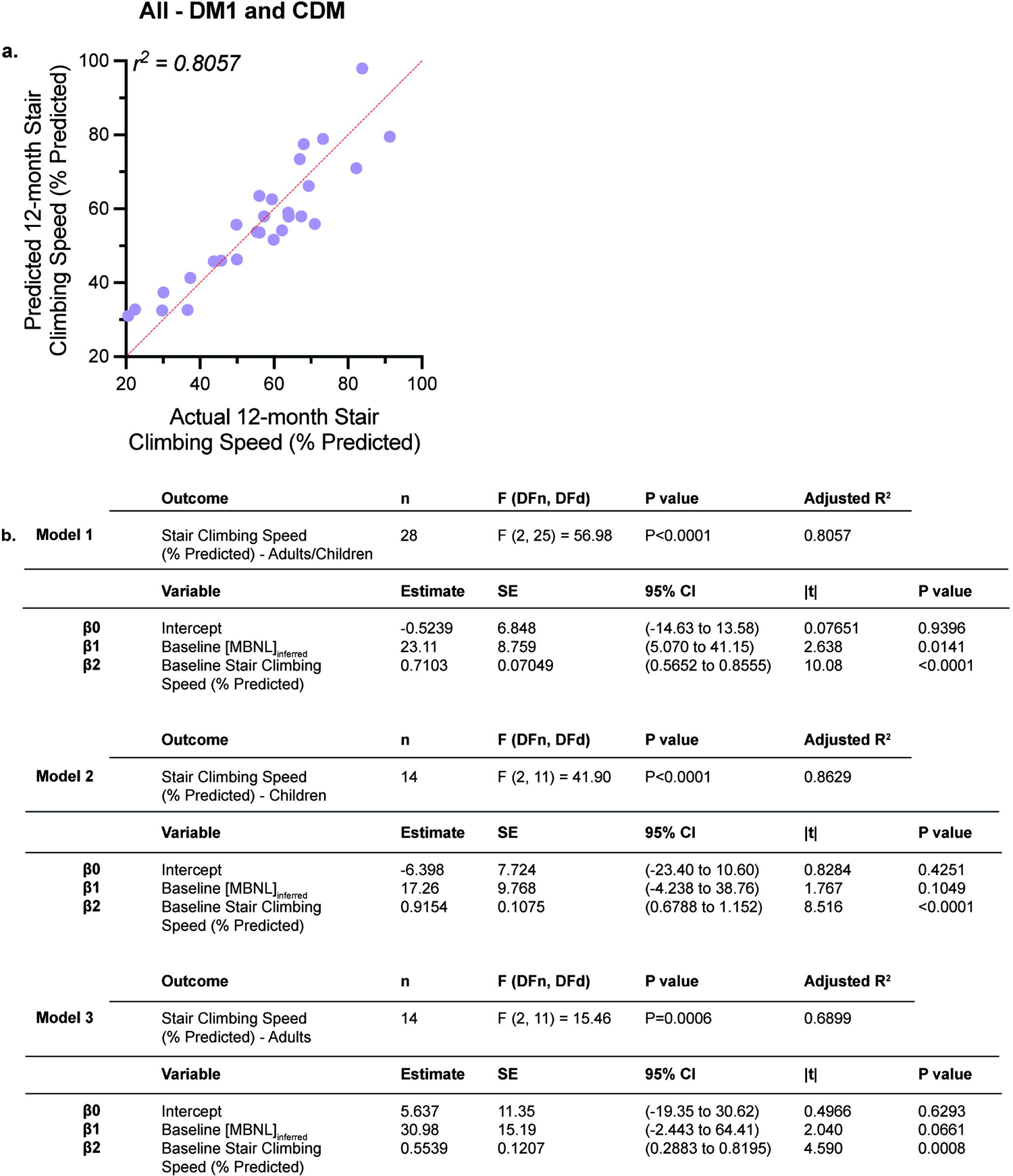
Multiple linear regression predictive modeling of 12-month stair climbing velocity in DM1 participants. (a.) Multiple linear regression plot showing observed 12-month stair climbing velocity (% predicted) versus predicted 12-month stair climbing velocity (% predicted) for all DM1 individuals (adults and children). (b.) Intercept table of multiple linear regression models for complete DM1 cohort (Model 1), CDM cohort alone (Model 2), and adult cohort alone (Model 3). Multiple statistical elements are reported (F = F-distribution (DFn = degrees of freedom numerator, DFd = degrees of freedom denominator), SE = standard error, 95% CI = 95% confidence interval, |t| = t-test statistic). Extended intercept tables for all three models are provided in Supplemental Table 7.

## Discussion

In this study we have identified associations between physical function and RNA mis-splicing in skeletal muscle as captured by [MBNL]_inferred_, a composite metric of global alternative splicing dysregulation, in adults and children with DM1. This study describes significant correlations between mis-splicing and timepoint-matched assessments of clinical performance in CDM children. Despite the availability of a small cohort of children and the wide age range assessed, the correlations observed across outcome assessments are surprisingly strong, underscoring the potential associations at play between alternative splicing dysregulation and phenotypic metrics shared between both DM1 adults and CDM children (Figures 2-4).

Importantly, the regression model equations developed above indicate that there are pathways forward in being able to predict levels of disease-associated spliceopathy in skeletal muscle within DM1 individuals using clinical outcome measures alone (Figure 5). Given the variable degree of mis-splicing in CDM, the ability to detect those children optimally suited for clinical trials by clinical outcome assessments may lessen the need for muscle biopsies in the future. The 12-month predictive multiple linear regression modeling analyses using baseline measures of alternative splicing dysregulation and baseline functional outcome performance points to the prognostic utility of measures such as [MBNL]_inferred_ (Figures 6-7). [MBNL]_inferred_ consistently trended towards significance in the 12-month predictive models for adults with DM1; the small sample sizes (n≤ 15) likely contributed to the lack of statistical significance observed.

Contrary to adults, it appears that [MBNL]_inferred_ is unlikely to contribute to 12-month predictive models in children with CDM, likely due to the extensive dynamic and variable pattern of alternative splicing changes that are occurring throughout development (Figure 1). This is critical to keep in mind as we move forward in designing clinical trials for children with CDM, as alternative splicing dysregulation in this patient population is dynamic and might confound results depending on age at assessment and trajectory of CDM disease progression (CDM_infant_ vs. CDM_child_ vs. CDM_adolescent_).

Overall, there was a positive correlation between [MBNL]_inferred_ and motor/strength performance in all individuals with DM1 regardless of age of onset. Individuals with lower [MBNL]_inferred_ levels reflective of extensive dysregulated RNA splicing were not able to walk as far, run as fast, and took longer to climb stairs. In children with CDM, those with lower [MBNL]_inferred_ levels also exhibited reduced dexterity on the 9-hole peg test, although this pattern was not seen in DM1 adults. Given the severe phenotype in infancy, it is reasonable to expect that this severe motor impairment may allow for detection of change on the 9-hole peg test compared to the less severely affected adults with DM1. Most strikingly, individuals with higher [MBNL]_inferred_ levels performed more similarly to unaffected individuals as measured by quantitative muscle testing (knee extension, ankle dorsiflexors, grip strength). Importantly, these associations were not substantially impacted by choice of the muscle biopsied or QMT testing method; [MBNL]_inferred_ was derived from vastus lateralis biopsies in children with CDM whereas distal tibialis anterior biopsies were utilized for adult-onset DM1 participants. Despite this difference in muscle groups used to measure RNA mis-splicing, CDM children alone had a similar magnitude, if not stronger, association between [MBNL]_inferred_ and measures of distal strength (hand grip and ankle dorsiflexion) when compared to our analyses that included all DM1 participants. Altogether, the results suggest that these associations are more indicative of global alterations in muscle function and strength rather than these patterns being reflective of one specific muscle group.

Perhaps the most surprising finding of these analyses was the observed lack of correlation between [MBNL]_inferred_ or *CLCN1*/*CACNA1S* mis-splicing levels with quantitative measures of myotonia in CDM children. Previous work has shown that increased inclusion of *CLCN1* exon 7a leads to reduced density of CLCN1 within muscle fibers and myotonic discharges. Correction of this mis-splicing via targeted antisense oligonucleotides is sufficient to rescue myotonia in DM1 mouse models^46^. Mis-splicing of the CaV1.1 calcium channel encoded by *CACNA1S* has also been linked with exacerbated myotonia in these model systems (ref). Consistent with these reports, increased inclusion of *CLCN1* exon7a or exclusion of *CACNA1S* exon29 was associated moderately with longer vHOT_thumb_ times in all DM1 participants; no such associations of myotonia with causative RNA mis-splicing events have been previously reported in human DM1 studies. However, these associations did not hold true in CDM children alone - all CDM participants, regardless of sub-cohort classification, [MBNL]_inferred_, or *CLCN1*/*CACNA1S* PSI values, had low vHOT_thumb_ times (< 3.2 s). In fact, CDM_adolescent_ individuals who had nearly 100% *CLCN1* exon inclusion, like the most severely affected DM1 adults, possessed minimal quantitative myotonia. Similar patterns were observed for *CACNA1S* mis-splicing. Altogether, these data suggest that while patterns of DM spliceopathy previously linked to myotonia are preserved in CDM skeletal muscle, these pediatric patients are not experiencing clinical myotonia in the same manner as DM1 adults. Future work will be required to evaluate the molecular mechanisms behind this lack of observable myotonia in the CDM population. In total, these data suggest that myotonia measures appear to be a poor outcome assessment for children with CDM, especially as correction of myotonia and the associated RNA splicing patterns have been proposed as early markers of therapeutic target engagement in DM1 clinical trials.

Previous work by our group has shown that CDM children display a triphasic pattern of disease progression that is mirrored by changes in RNA mis-splicing^13,28^. While the CDM participants evaluated here were sampled cross-sectionally, the results captured in this study begin to verify that the triphasic pattern of alternative splicing dysregulation in CDM matches the observed changes in physical function. However, we were unable to fully validate this proposed pattern given the limited sample size and data availability within select CDM sub-cohorts, notably CDM infants (< 2 years). Few clinical measures were able to be assessed for these individuals due to the developmental constraints of this age group. Future directions include improving our clinical outcome measures to include infant-specific strength/motor testing. Another challenge in obtaining larger sample sizes lies in the difficulty in obtaining reliable outcomes from patients with physical, cognitive, and behavioral impairments. Due to these challenges, it is unclear whether clinical outcome measures reflect true maximum physical ability, or rather a child’s ability to understand and follow instructions. This could be especially relevant in measures such as the 9-hole peg test. Future studies with expanded cohort sizes will need to be done to assess the impacts of cognitive impairment as a potential confounding variable in physical outcome measure performance.

The development of disease-modifying therapies for adults with DM1 has rapidly accelerated in recent years with many in early-phase clinical trials. Given the shared genetic basis of disease, there is significant interest in extending these trials into the CDM population. However, a major limitation of these efforts has been insufficient understanding of the natural history of disease progression in children and how the molecular pathogenesis, particularly alternative splicing dysregulation, correlates with clinical outcome measures. This analysis connects measures of splicing dysregulation to a range of clinical outcome measures in a cross-sectional cohort including both adult DM1 and CDM participants.

Furthermore, the analyses here provide the foundation for determination of high-performing clinical outcome assessments (COAs) for use in CDM clinical trials in older pediatric patients (> 6 years). This work, in combination with our characterization of CDM spliceopathy across pediatric development^28^, is vital in identifying the effective timing of therapeutic administration to CDM children for clinical trial success whereby therapeutic benefit outpaces the natural dynamic changes in RNA mis-splicing, especially given the rapid improvement observed in the first years of life (Figure 1). The regression models developed in this study may assist in the refinement of clinical trial design and offer a non-invasive methodology for prediction of spliceopathy using COAs alone when screening for trial inclusion. They may also reduce reliance on muscle biopsies to assess target engagement of therapeutic agents. The strength of these predictive equations can be improved through replication and validation in a secondary DM1 cohort. Additionally, these predicative equations are based on inferences of free [MBNL] in skeletal muscle and may not be reflective of disease severity and clinical performance for non-musculoskeletal metrics. Overall, these results provide the framework for both the utility of a muscle-based biomarker and clinical trial design in children with CDM.

## Supporting information

DMCRN Consortium Members

## Acknowledgments

Sources of Support: This study is supported by NINDS (R01NS104010), FDA (7R01FD006071), Muscular Dystrophy Association, Myotonic Dystrophy Foundation, Novartis, Dyne, Avidity, PepGen, and Takeda.

## Author Contributions

Conception and Design of Study: JMH, MAH, NEJ

Acquisition and Analysis of Data: JMH, MP, KB, KI, AB, AJ, KB, JD, MM, JB, KC, JB, MK, NEJ, MAH

Drafting a Significant Portion of Manuscript: JMH, MK, NEJ, MAH

Please see supplementary file labeled “DMCRN Consortium Members.”

## Data Availability

RNA sequencing data used to define [MBNL]inferred are available in Sequence Read Archive at https://www.ncbi.nlm.nih.gov/sra, Accession [PRJNA1079722] and [PRJNA830511].

