## Supplemental Table & Figure Legends for "RNA mis-splicing in children with myotonic dystrophy is associated with physical function"

**Supplemental Figure 1:** Myotonia measures in adult DM1 and CDM participants correlate disparately with specific skeletal muscle mis-spliced events. (a.) Correlation between CACNA1S exon 29 percent spliced in (PSI) and vHOT_thumb_ times in all DM1 individuals and CDM individuals. Correlations were calculated using a two-tailed Spearman test.

**Supplementary Table 1:** Demographic information of participants included in this study, including age at biopsy, visit number, sex, CTG repeat expansion size, [MBNL]_inferred_, and CDM sub-cohort classification, where applicable. Visit number is indicative of the study visit, where 0 = baseline visit, 1 = 3-month visit, and 2 = 12-month visit. Some sampled individuals had CTG repeat lengths reported as a range and thus the lowest value of the range was used as their reported repeat length. CDM individuals with repeat biopsy are indicated with * (CDM-01), ** (CDM-30), and *** (CDM-37), respectively. Missing values are left blank in the table.

**Supplementary Table 2:** Raw values for clinical outcome measures associated with each participant ID. CDM individuals with repeat biopsy are indicated with * (CDM-01), ** (CDM-30), and *** (CDM-37), respectively. Missing values are left blank in the table. s = seconds, m = meters, kgf = kilogram-force.

**Supplementary Table 3:** *Complete intercept table of multiple linear regression Model 1 for estimating [MBNL]_inferred_ in all participants (adults and children).* Companion table to regression model reported in Figure 5 with all elements and associated statistical tests provided.

**Supplementary Table 4:** *Complete intercept table of multiple linear regression Model 2 for estimating [MBNL]_inferred_ in CDM participants.* Companion table to regression model reported in Figure 5 with all elements and associated statistical tests provided.

**Supplementary Table 5:** *Complete intercept tables of multiple linear regression for estimating 12-month walk/running velocity (% predicted) in adults and children.* Companion table to regression models reported in Figure 6 with all elements and associated statistical tests provided.

**Supplementary Table 6:** *Complete intercept tables of multiple linear regression for estimating 12-month ankle dorsiflexion strength (% predicted) in adults and children.* Companion table to regression models reported in Figure 6 with all elements and associated statistical tests provided.

**Supplementary Table 7:** *Complete intercept tables of multiple linear regression for estimating 12-month stair climbing speed (% predicted) in adults and children.* Companion table to regression models reported in Figure 7 with all elements and associated statistical tests provided.
