## Supplemental Table 1 for "RNA mis-splicing in children with myotonic dystrophy is associated with physical function"

| **Sample ID** | **Biopsy Location** | **Cohort** | **CDM Sub-cohort** | **Visit** | **Sex** | **Age at Biopsy (yrs.)** | **CTG Repeat** | **[MBNL]_inferred_** |
| --- | --- | --- | --- | --- | --- | --- | --- | --- |
| CDM-01* | vastus lateralis | CDM | CDM infant | 0 | M | 0.04 | 1500 | 0.015 |
| CDM-02 | vastus lateralis | CDM | CDM infant | 0 | M | 0.125 |  | 0.013 |
| CDM-03 | vastus lateralis | CDM | CDM infant | 0 | M | 0.25 |  | 0.021 |
| CDM-04 | vastus lateralis | CDM | CDM infant | 0 | F | 1 | 1700 | 0.139 |
| CDM-05 | vastus lateralis | CDM | CDM infant | 0 | M | 1 |  | 0.681 |
| CDM-06 | vastus lateralis | CDM | CDM child | 0 | M | 2 |  | 0.83 |
| CDM-07 | vastus lateralis | CDM | CDM infant | 0 | M | 0.92 |  | 0.349 |
| CDM-08 | vastus lateralis | CDM | CDM child | 0 | M | 7 |  | 0.441 |
| CDM-09 | vastus lateralis | CDM | CDM child | 0 | M | 3 |  | 0.907 |
| CDM-10 | vastus lateralis | CDM | CDM child | 0 | F | 2.5 | 1500 | 0.522 |
| CDM-11 | vastus lateralis | CDM | CDM child | 0 | M | 2.5 | 750 | 0.778 |
| CDM-12 | vastus lateralis | CDM | CDM child | 0 | M | 5 | 1000 | 0.493 |
| CDM-13 | vastus lateralis | CDM | CDM child | 0 | M | 5 | 1060 | 0.519 |
| CDM-14 | vastus lateralis | CDM | CDM child | 0 | M | 5 | 938 | 0.643 |
| CDM-15 | vastus lateralis | CDM | CDM child | 0 | M | 5 | 1000 | 0.674 |
| CDM-16 | vastus lateralis | CDM | CDM child | 0 | F | 5 | 550 | 0.92 |
| CDM-17 | vastus lateralis | CDM | CDM child | 0 | F | 6 | 1450 | 0.757 |
| CDM-18 | vastus lateralis | CDM | CDM child | 0 | M | 6 | 1300 | 0.536 |
| CDM-19 | vastus lateralis | CDM | CDM child | 0 | M | 6 | 465 | 0.916 |
| CDM-20 | vastus lateralis | CDM | CDM child | 0 | M | 6 | 1505 | 0.715 |
| CDM-21 | vastus lateralis | CDM | CDM child | 0 | M | 7 | 950 | 0.826 |
| CDM-22 | vastus lateralis | CDM | CDM adolescent-1 | 0 | M | 8 | 986 | 0.702 |
| CDM-23* | vastus lateralis | CDM | CDM adolescent-1 | 0 | M | 8 | 1500 | 0.577 |
| CDM-24 | vastus lateralis | CDM | CDM adolescent-1 | 0 | F | 8 | 773 | 0.903 |
| CDM-25 | vastus lateralis | CDM | CDM adolescent-2 | 0 | M | 8 | 1500 | 0.214 |
| CDM-26 | vastus lateralis | CDM | CDM adolescent-2 | 0 | F | 8 | 1473 | 0.264 |
| CDM-27 | vastus lateralis | CDM | CDM adolescent-1 | 0 | F | 8 | 1800 | 0.729 |
| CDM-28 | vastus lateralis | CDM | CDM child | 0 | F | 7 | 983 | 0.712 |
| CDM-29 | vastus lateralis | CDM | CDM adolescent-1 | 0 | F | 8 |  | 0.608 |
| CDM-30** | vastus lateralis | CDM | CDM adolescent-1 | 0 | M | 9 | 854 | 0.933 |
| CDM-31 | vastus lateralis | CDM | CDM adolescent-2 | 0 | F | 10 |  | 0.306 |
| CDM-32 | vastus lateralis | CDM | CDM adolescent-1 | 0 | M | 10 | 1200 | 0.824 |
| CDM-33 | vastus lateralis | CDM | CDM adolescent-1 | 0 | M | 11 | 1050 | 0.787 |
| CDM-34 | vastus lateralis | CDM | CDM adolescent-1 | 0 | F | 11 | 1136 | 0.544 |
| CDM-35 | soleus | CDM | CDM adolescent-2 | 0 | F | 11 |  | 0.151 |
| CDM-36 | vastus lateralis | CDM | CDM adolescent-2 | 0 | F | 12 | 2530 | 0.275 |
| CDM-37*** | vastus lateralis | CDM | CDM adolescent-1 | 0 | F | 12 | 710 | 0.727 |
| CDM-38 | vastus lateralis | CDM | CDM adolescent-2 | 0 | F | 14 | 1480 | 0.018 |
| CDM-39 | vastus lateralis | CDM | CDM adolescent-1 | 0 | M | 14 | 450 | 0.863 |
| CDM-40 | vastus lateralis | CDM | CDM adolescent-2 | 0 | F | 16 |  | 0.352 |
| CDM-41*** | vastus lateralis | CDM | CDM adolescent-1 | 0 | F | 16 | 710 | 0.748 |
| CDM-42 | vastus lateralis | CDM | CDM adolescent-1 | 0 | M | 16 |  | 0.903 |
| CDM-43** | vastus lateralis | CDM | CDM adolescent-1 | 0 | M | 13 |  | 0.8 |
| CDM-44 |  | CDM | CDM infant | 2 | M | 1 | 1500 |  |
| CDM-45 |  | CDM | CDM infant | 2 | M | 1 |  |  |
| CDM-46 |  | CDM | CDM infant | 2 | M | 1 |  |  |
| CDM-47 |  | CDM | CDM infant | 2 | F | 2 | 1700 |  |
| CDM-48 |  | CDM | CDM child | 2 | F | 3 | 1500 |  |
| CDM-49 |  | CDM | CDM child | 2 | M | 4 |  |  |
| CDM-50 |  | CDM | CDM child | 2 | M | 6 | 1000 |  |
| CDM-51 |  | CDM | CDM child | 2 | M | 6 | 1060 |  |
| CDM-52 |  | CDM | CDM child | 2 | M | 6 | 938 |  |
| CDM-53 |  | CDM | CDM child | 2 | M | 6 | 1000 |  |
| CDM-54 |  | CDM | CDM child | 2 | F | 6 | 550 |  |
| CDM-55 |  | CDM | CDM child | 2 | F | 7 | 1450 |  |
| CDM-56 |  | CDM | CDM child | 2 | M | 7 | 1300 |  |
| CDM-57 |  | CDM | CDM child | 2 | M | 7 | 465 |  |
| CDM-58 |  | CDM | CDM child | 2 | M | 7 | 1505 |  |
| CDM-59 |  | CDM | CDM child | 2 | M | 8 | 950 |  |
| CDM-60 |  | CDM | CDM child | 2 | M | 9 | 986 |  |
| CDM-61 |  | CDM | CDM adolescent-1 | 2 | M | 9 | 1500 |  |
| CDM-62 |  | CDM | CDM adolescent-1 | 2 | F | 9 | 773 |  |
| CDM-63 |  | CDM | CDM adolescent-2 | 2 | M | 9 | 1500 |  |
| CDM-64 |  | CDM | CDM adolescent-2 | 2 | F | 9 | 1473 |  |
| CDM-65 |  | CDM | CDM adolescent-1 | 2 | F | 9 | 1800 |  |
| CDM-66 |  | CDM | CDM adolescent-1 | 2 | F | 8 | 983 |  |
| CDM-67 |  | CDM | CDM adolescent-1 | 2 | F | 9 |  |  |
| CDM-68 |  | CDM | CDM adolescent-1 | 2 | M | 10 | 854 |  |
| CDM-69 |  | CDM | CDM adolescent-2 | 2 | F | 11 |  |  |
| CDM-70 |  | CDM | CDM adolescent-1 | 2 | M | 11 | 1200 |  |
| CDM-71 |  | CDM | CDM adolescent-1 | 2 | M | 12 | 1050 |  |
| CDM-72 |  | CDM | CDM adolescent-1 | 2 | F | 12 | 1136 |  |
| CDM-73 |  | CDM | CDM adolescent-2 | 2 | F | 12 |  |  |
| CDM-74 |  | CDM | CDM adolescent-2 | 2 | F | 13 | 2530 |  |
| CDM-75 |  | CDM | CDM adolescent-1 | 2 | F | 13 | 710 |  |
| CDM-76 |  | CDM | CDM adolescent-2 | 2 | F | 15 | 1480 |  |
| CDM-77 |  | CDM | CDM adolescent-1 | 2 | M | 15 | 450 |  |
| DM1-01 | tibialis anterior | DM1 |  | 0 | F | 34 | 340 | 0.58 |
| DM1-02 | tibialis anterior | DM1 |  | 0 | F | 31 | 350 | 0.563 |
| DM1-03 | tibialis anterior | DM1 |  | 0 | M | 43 |  | 0.231 |
| DM1-04 | tibialis anterior | DM1 |  | 0 | M | 37 | 350 | 0.507 |
| DM1-05 | tibialis anterior | DM1 |  | 0 | F | 33 |  | 0.386 |
| DM1-06 | tibialis anterior | DM1 |  | 0 | M | 28 |  | 0.634 |
| DM1-07 | tibialis anterior | DM1 |  | 0 | F | 30 |  | 0.446 |
| DM1-08 | tibialis anterior | DM1 |  | 0 | F | 43 |  | 0.036 |
| DM1-09 | tibialis anterior | DM1 |  | 0 | M | 28 | 505 | 0.221 |
| DM1-10 | tibialis anterior | DM1 |  | 0 | M | 43 | 893 | 0.494 |
| DM1-11 | tibialis anterior | DM1 |  | 0 | F | 39 | 600 | 0.303 |
| DM1-12 | tibialis anterior | DM1 |  | 0 | M | 41 | 866 | 0.364 |
| DM1-13 | tibialis anterior | DM1 |  | 0 | M | 41 | 746 | 0.38 |
| DM1-14 | tibialis anterior | DM1 |  | 0 | F | 54 |  | 0.133 |
| DM1-15 | tibialis anterior | DM1 |  | 0 | M | 49 |  | 0.33 |
| DM1-16 | tibialis anterior | DM1 |  | 0 | F | 56 |  | 0.456 |
| DM1-17 | tibialis anterior | DM1 |  | 0 | F | 57 | 179 | 0.081 |
| DM1-18 | tibialis anterior | DM1 |  | 0 | M | 36 | 677 | 0.208 |
| DM1-19 | tibialis anterior | DM1 |  | 0 | M | 38 | 720 | 0.032 |
| DM1-20 | tibialis anterior | DM1 |  | 0 | M | 35 |  | 0.025 |
| DM1-21 | tibialis anterior | DM1 |  | 0 | M | 41 | 441 | 0.054 |
| DM1-22 | tibialis anterior | DM1 |  | 1 | F | 35 | 340 | 0.595 |
| DM1-23 | tibialis anterior | DM1 |  | 1 | M | 44 | 350 | 0.437 |
| DM1-24 | tibialis anterior | DM1 |  | 1 | M | 29 |  | 0.478 |
| DM1-25 | tibialis anterior | DM1 |  | 1 | F | 30 |  | 0.416 |
| DM1-26 | tibialis anterior | DM1 |  | 1 | F | 44 |  | 0.071 |
| DM1-27 | tibialis anterior | DM1 |  | 1 | M | 29 | 505 | 0.197 |
| DM1-28 | tibialis anterior | DM1 |  | 1 | F | 39 | 600 | 0.015 |
| DM1-29 | tibialis anterior | DM1 |  | 1 | M | 41 | 866 | 0.219 |
| DM1-30 | tibialis anterior | DM1 |  | 1 | F | 55 |  | 0.291 |
| DM1-31 | tibialis anterior | DM1 |  | 1 | M | 49 |  | 0.038 |
| DM1-32 | tibialis anterior | DM1 |  | 1 | F | 57 | 179 | 0.128 |
| DM1-33 | tibialis anterior | DM1 |  | 1 | M | 37 | 677 | 0.092 |
| DM1-34 | tibialis anterior | DM1 |  | 1 | M | 38 | 720 | 0.052 |
| DM1-35 | tibialis anterior | DM1 |  | 1 | M | 36 |  | 0.042 |
| DM1-36 | tibialis anterior | DM1 |  | 1 | M | 42 | 441 | 0.034 |
| DM1-37 | tibialis anterior | DM1 |  | 0 | F | 31 |  | 0.83 |
| DM1-38 | tibialis anterior | DM1 |  | 0 | M | 35 |  | 0.825 |
| DM1-39 | tibialis anterior | DM1 |  | 0 | F | 42 |  | 0.537 |
| DM1-40 | tibialis anterior | DM1 |  | 0 | F | 42 | 313 | 0.488 |
| DM1-41 | tibialis anterior | DM1 |  | 0 | F | 41 | 133 | 0.638 |
| DM1-42 | tibialis anterior | DM1 |  | 0 | F | 45 |  | 0.672 |
| DM1-43 | tibialis anterior | DM1 |  | 0 | F | 33 | 170 | 0.511 |
| DM1-44 | tibialis anterior | DM1 |  | 0 | F | 38 |  | 0.503 |
| DM1-45 | tibialis anterior | DM1 |  | 0 | F | 38 | 300 | 0.627 |
| DM1-46 | tibialis anterior | DM1 |  | 0 | M | 45 | 108 | 0.956 |
| DM1-47 | tibialis anterior | DM1 |  | 0 | F | 21 |  | 0.749 |
| DM1-48 | tibialis anterior | DM1 |  | 0 | M | 20 |  | 0.517 |
| DM1-49 | tibialis anterior | DM1 |  | 0 | F | 42 |  | 0.56 |
| DM1-50 | tibialis anterior | DM1 |  | 0 | F | 62 | 336 | 0.479 |
| DM1-51 | tibialis anterior | DM1 |  | 0 | F | 44 |  | 0.392 |
| DM1-52 | tibialis anterior | DM1 |  | 0 | F | 32 | 750 | 0.425 |
| DM1-53 | tibialis anterior | DM1 |  | 0 | M | 31 |  | 0.748 |
| DM1-54 | tibialis anterior | DM1 |  | 0 | F | 21 | 300 | 0.729 |
| DM1-55 | tibialis anterior | DM1 |  | 0 | F | 44 |  | 0.874 |
| DM1-56 | tibialis anterior | DM1 |  | 0 | F | 38 | 200 | 0.491 |
| DM1-57 | tibialis anterior | DM1 |  | 0 | M | 69 | 98 | 0.772 |
| DM1-58 | tibialis anterior | DM1 |  | 1 | F | 42 |  | 0.604 |
| DM1-59 | tibialis anterior | DM1 |  | 1 | F | 42 | 313 | 0.098 |
| DM1-60 | tibialis anterior | DM1 |  | 1 | F | 41 | 133 | 0.651 |
| DM1-61 | tibialis anterior | DM1 |  | 1 | F | 45 |  | 0.928 |
| DM1-62 | tibialis anterior | DM1 |  | 1 | F | 38 |  | 0.289 |
| DM1-63 | tibialis anterior | DM1 |  | 1 | F | 38 | 300 | 0.358 |
| DM1-64 | tibialis anterior | DM1 |  | 1 | M | 45 | 108 | 0.797 |
| DM1-65 | tibialis anterior | DM1 |  | 1 | F | 21 |  | 0.563 |
| DM1-66 | tibialis anterior | DM1 |  | 1 | M | 20 |  | 0.368 |
| DM1-67 | tibialis anterior | DM1 |  | 1 | F | 62 | 336 | 0.412 |
| DM1-68 | tibialis anterior | DM1 |  | 1 | F | 32 | 750 | 0.433 |
| DM1-69 | tibialis anterior | DM1 |  | 1 | M | 31 |  | 0.784 |
| DM1-70 | tibialis anterior | DM1 |  | 1 | F | 21 | 300 | 0.707 |
| DM1-71 | tibialis anterior | DM1 |  | 1 | F | 44 |  | 0.819 |
| DM1-72 | tibialis anterior | DM1 |  | 1 | F | 38 | 200 | 0.412 |
| DM1-73 | tibialis anterior | DM1 |  | 1 | M | 69 | 98 | 0.722 |
| DM1-74 |  | DM1 |  | 2 | F | 43 |  |  |
| DM1-75 |  | DM1 |  | 2 | F | 43 | 313 |  |
| DM1-76 |  | DM1 |  | 2 | F | 42 | 133 |  |
| DM1-77 |  | DM1 |  | 2 | F | 34 |  |  |
| DM1-78 |  | DM1 |  | 2 | F | 39 | 108 |  |
| DM1-79 |  | DM1 |  | 2 | M | 46 |  |  |
| DM1-80 |  | DM1 |  | 2 | F | 22 |  |  |
| DM1-81 |  | DM1 |  | 2 | M | 21 | 336 |  |
| DM1-82 |  | DM1 |  | 2 | F | 63 |  |  |
| DM1-83 |  | DM1 |  | 2 | F | 45 | 300 |  |
| DM1-84 |  | DM1 |  | 2 | F | 33 |  |  |
| DM1-85 |  | DM1 |  | 2 | M | 32 | 200 |  |
| DM1-86 |  | DM1 |  | 2 | F | 22 | 98 |  |
| DM1-87 |  | DM1 |  | 2 | F | 39 |  |  |
| DM1-88 |  | DM1 |  | 2 | M | 70 |  |  |
