## Supplemental Table 2 for "RNA mis-splicing in children with myotonic dystrophy is associated with physical function"

| **Sample ID** | **Myotonia Scale** | **Myotonia vHOT_thumb_ (s)** | **9-Hole Peg Test (s)** | **6-Minute Walk (m)** | **Stair Climbing Velocity (stair/s)** | **Walk/Run Velocity (m/sec)** | **Knee Extension (kgf)** | **Grip Strength (kgf)** | **Ankle Dorsiflexion (kgf)** |
| --- | --- | --- | --- | --- | --- | --- | --- | --- | --- |
| CDM-01* |  |  |  |  |  |  |  |  |  |
| CDM-02 |  |  |  |  |  |  |  |  |  |
| CDM-03 |  |  |  |  |  |  |  |  |  |
| CDM-04 | 1.00 |  |  |  |  |  |  |  |  |
| CDM-05 |  |  |  |  |  |  |  |  |  |
| CDM-06 |  |  |  |  |  |  |  |  |  |
| CDM-07 |  |  |  |  |  |  |  |  |  |
| CDM-08 |  |  |  |  |  |  |  |  |  |
| CDM-09 |  |  |  |  |  |  |  |  |  |
| CDM-10 | 1.00 |  |  |  |  |  |  |  |  |
| CDM-11 | 1.00 |  |  |  |  |  |  |  |  |
| CDM-12 | 1.00 |  | 50.05 | 248.00 | 0.74 | 1.12 |  | 1.80 |  |
| CDM-13 | 1.00 |  | 75.50 |  | 1.18 | 1.31 |  | 2.63 |  |
| CDM-14 | 1.00 |  | 47.90 | 325.00 | 1.25 | 1.94 |  | 3.30 |  |
| CDM-15 | 4.00 |  |  |  | 1.24 | 1.73 |  | 3.85 |  |
| CDM-16 | 1.00 |  | 38.83 | 375.00 | 1.67 | 2.33 | 6.93 | 4.25 | 3.92 |
| CDM-17 | 3.00 | 0.48 | 30.59 | 352.00 | 1.05 | 1.90 | 5.67 | 3.87 |  |
| CDM-18 |  |  |  | 225.00 | 0.65 | 1.56 |  |  |  |
| CDM-19 | 3.00 | 0.65 |  | 450.00 | 1.27 | 2.98 | 7.50 | 5.17 | 2.82 |
| CDM-20 | 1.00 |  | 61.68 | 227.00 | 0.56 | 1.14 |  | 4.10 |  |
| CDM-21 | 5.00 | 0.26 | 37.49 | 473.00 | 2.03 | 2.72 | 12.43 | 7.52 | 8.28 |
| CDM-22 | 3.00 | 0.37 | 37.21 | 399.00 | 0.67 | 1.30 | 7.45 | 5.22 |  |
| CDM-23* | 5.00 | 0.51 | 77.00 | 300.00 | 0.31 | 1.11 | 6.70 | 4.20 |  |
| CDM-24 | 4.00 | 3.10 | 27.22 | 567.00 | 1.68 | 2.86 | 13.73 | 9.67 | 9.62 |
| CDM-25 | 1.00 |  |  | 294.00 | 1.11 | 2.17 |  |  | 2.75 |
| CDM-26 | 1.00 | 0.41 |  |  |  |  | 3.82 | 2.40 | 1.60 |
| CDM-27 | 1.00 |  |  | 176.00 | 0.33 | 0.79 |  | 2.43 |  |
| CDM-28 | 3.00 |  | 39.51 | 364.00 | 1.11 | 2.08 | 8.33 | 4.77 |  |
| CDM-29 | 6.00 |  |  | 283.00 | 1.18 | 2.17 |  |  |  |
| CDM-30** | 1.00 |  | 31.10 | 460.00 | 1.74 | 2.61 | 9.63 | 8.65 | 5.60 |
| CDM-31 | 1.00 | 0.31 | 122.33 | 102.00 | 0.14 |  |  | 3.07 |  |
| CDM-32 | 2.00 | 3.20 | 31.70 |  | 1.76 | 2.27 | 11.87 | 4.70 |  |
| CDM-33 | 2.00 |  |  | 304.00 | 0.72 | 1.58 | 4.72 | 4.88 |  |
| CDM-34 | 2.00 | 1.05 | 27.42 | 322.00 | 1.85 | 2.43 | 11.57 | 5.60 | 5.02 |
| CDM-35 | 1.00 |  |  | 416.00 | 1.86 | 2.33 |  | 11.90 | 2.32 |
| CDM-36 | 3.00 | 0.98 | 48.36 | 390.00 | 0.68 | 1.60 | 7.45 | 8.37 |  |
| CDM-37*** | 3.00 | 0.55 | 21.13 | 590.00 | 2.33 | 2.79 | 12.78 | 7.70 | 9.52 |
| CDM-38 | 2.00 | 1.55 | 22.27 | 466.00 | 1.58 | 2.22 | 11.20 | 11.72 | 4.70 |
| CDM-39 | 2.00 | 0.30 | 22.79 | 513.00 | 2.17 | 3.43 | 16.23 | 21.27 | 11.30 |
| CDM-40 |  |  |  |  |  |  |  |  |  |
| CDM-41*** |  |  |  |  |  |  |  |  |  |
| CDM-42 |  |  |  |  |  |  |  |  |  |
| CDM-43** |  |  |  |  |  |  |  |  |  |
| CDM-44 |  |  |  |  |  |  |  |  |  |
| CDM-45 |  |  |  |  |  |  |  |  |  |
| CDM-46 |  |  |  |  |  |  |  |  |  |
| CDM-47 |  |  |  |  |  |  |  |  |  |
| CDM-48 |  |  |  |  |  |  |  |  |  |
| CDM-49 | 1.00 | 1.00 | 43.50 | 278.00 | 1.37 | 1.88 | 6.38 | 3.67 |  |
| CDM-50 | 1.00 |  | 75.94 | 313.00 | 0.73 | 1.17 |  | 4.50 |  |
| CDM-51 | 4.00 |  | 55.30 | 253.00 | 1.34 | 1.40 |  | 2.18 |  |
| CDM-52 | 6.00 |  | 69.67 |  | 1.54 | 1.79 |  | 3.57 |  |
| CDM-53 |  |  |  |  |  |  |  |  |  |
| CDM-54 |  |  |  |  |  |  |  |  |  |
| CDM-55 | 2.00 |  | 33.15 | 309.00 | 1.31 | 2.14 | 7.80 | 4.62 | 7.80 |
| CDM-56 |  |  |  | 299.00 | 0.54 | 1.45 |  |  |  |
| CDM-57 |  |  | 36.79 | 506.00 | 1.51 | 2.45 | 8.25 | 4.42 | 8.25 |
| CDM-58 |  |  |  |  |  |  |  |  |  |
| CDM-59 | 3.00 | 0.35 | 39.73 | 486.00 | 1.88 | 2.75 | 10.47 | 6.48 | 10.47 |
| CDM-60 | 1.00 |  | 54.50 | 350.00 | 0.75 | 1.46 | 5.57 | 4.50 |  |
| CDM-61 |  |  |  |  |  |  |  |  |  |
| CDM-62 |  |  |  |  |  |  |  |  |  |
| CDM-63 | 3.00 | 1.14 | 28.90 | 353.00 | 1.23 | 1.99 |  | 6.57 |  |
| CDM-64 |  |  |  |  |  |  |  |  |  |
| CDM-65 |  |  |  |  |  |  |  |  |  |
| CDM-66 | 4.00 | 1.42 | 27.37 | 410.00 | 1.19 | 2.19 | 10.85 | 7.67 | 10.85 |
| CDM-67 |  |  |  |  |  |  |  |  |  |
| CDM-68 | 1.00 |  | 31.63 | 553.00 | 1.89 | 2.86 | 7.52 | 5.50 | 3.23 |
| CDM-69 |  |  |  |  |  |  |  |  |  |
| CDM-70 |  |  |  |  |  |  |  |  |  |
| CDM-71 |  |  | 38.40 | 314.00 | 0.99 | 1.74 | 8.43 | 6.00 | 8.43 |
| CDM-72 | 2.00 | 18.50 | 32.00 | 324.00 | 2.35 | 2.55 | 5.52 | 4.65 | 5.52 |
| CDM-73 |  |  |  |  |  |  |  |  |  |
| CDM-74 |  |  |  |  |  |  |  |  |  |
| CDM-75 | 2.00 | 1.60 | 22.16 | 601.00 | 2.33 | 3.24 | 13.50 | 9.63 | 13.50 |
| CDM-76 | 2.00 | 2.67 | 23.68 | 467.00 | 1.46 | 2.54 | 11.75 | 12.10 | 11.75 |
| CDM-77 |  |  |  |  |  | 2.22 |  |  |  |
| DM1-01 | 3.00 | 7.17 |  | 567.00 |  | 1.98 | 20.53 | 19.66 | 11.00 |
| DM1-02 | 2.00 | 3.39 |  | 480.00 | 1.47 | 1.74 | 34.67 | 10.34 | 3.92 |
| DM1-03 | 5.00 | 5.21 |  | 321.50 | 1.53 | 1.36 | 17.89 | 11.16 | 2.93 |
| DM1-04 | 4.00 | 8.55 |  | 553.00 | 1.58 | 1.65 | 20.79 | 27.00 | 8.90 |
| DM1-05 | 5.00 | 5.67 |  | 443.00 | 1.06 | 0.69 | 26.49 | 12.21 | 5.60 |
| DM1-06 | 5.00 | 36.96 |  | 489.00 | 1.47 | 1.53 | 24.10 | 12.14 | 9.93 |
| DM1-07 | 4.00 | 3.40 |  | 304.00 | 1.26 | 1.24 | 16.79 | 9.85 | 5.92 |
| DM1-08 | 3.00 | 4.01 |  | 300.00 | 1.26 | 1.08 | 13.62 | 2.90 |  |
| DM1-09 | 2.00 | 16.91 |  | 465.00 | 1.88 | 1.67 | 19.96 | 15.71 | 6.14 |
| DM1-10 | 2.00 | 5.00 |  | 358.00 | 1.42 | 1.29 | 19.92 | 11.35 | 5.32 |
| DM1-11 | 2.00 | 3.16 |  | 428.00 | 1.58 | 1.23 | 23.51 | 14.63 | 9.18 |
| DM1-12 | 5.00 | 6.04 |  | 470.00 | 1.62 | 1.51 | 14.36 | 9.43 | 4.67 |
| DM1-13 | 5.00 | 9.61 |  | 250.00 | 1.22 | 0.92 | 9.92 | 7.39 | 1.84 |
| DM1-14 | 3.00 | 5.23 |  | 303.00 | 0.82 | 1.09 | 9.89 | 7.98 | 2.07 |
| DM1-15 | 4.00 | 21.96 |  | 315.00 |  | 1.00 | 20.25 | 4.64 |  |
| DM1-16 | 2.00 | 3.37 |  | 150.00 | 0.50 |  | 17.04 | 13.27 | 4.58 |
| DM1-17 | 3.00 | 11.93 |  | 236.00 | 0.42 | 0.87 | 7.35 | 4.85 | 2.35 |
| DM1-18 | 3.00 | 20.65 |  | 371.00 | 1.53 | 1.27 | 16.53 | 6.56 | 6.61 |
| DM1-19 | 4.00 | 25.34 |  | 360.00 | 1.58 | 1.59 | 14.33 | 4.73 | 2.66 |
| DM1-20 | 2.00 |  |  |  | 1.12 | 1.28 | 42.88 | 7.63 | 3.76 |
| DM1-21 | 5.00 | 27.59 |  | 383.00 | 1.52 | 1.42 | 33.32 | 5.33 | 1.39 |
| DM1-22 | 4.00 | 3.65 |  | 545.00 | 2.25 | 1.75 | 12.34 | 17.11 | 11.00 |
| DM1-23 | 4.00 | 3.62 |  | 525.00 | 1.76 | 1.76 | 25.19 | 23.17 | 10.21 |
| DM1-24 | 5.00 | 35.92 |  | 473.00 | 1.58 | 1.53 | 27.07 | 11.94 | 9.39 |
| DM1-25 | 3.00 | 3.25 |  | 374.00 | 1.58 | 1.28 | 15.49 | 8.85 | 5.34 |
| DM1-26 | 1.00 | 4.53 |  | 306.00 | 1.24 | 0.89 | 12.74 | 3.16 |  |
| DM1-27 | 3.00 | 11.88 |  | 530.00 | 3.24 | 1.82 | 21.74 | 16.74 | 7.22 |
| DM1-28 | 4.00 | 6.63 |  | 408.00 | 1.65 | 1.19 | 34.49 | 14.78 | 11.40 |
| DM1-29 | 4.00 | 8.83 |  | 496.00 | 1.65 | 1.25 | 10.46 | 6.32 | 4.51 |
| DM1-30 | 2.00 | 8.96 |  | 309.00 | 1.01 | 1.03 | 7.77 | 8.35 | 2.43 |
| DM1-31 | 6.00 | 20.37 |  | 265.00 |  | 0.96 | 13.28 | 5.10 |  |
| DM1-32 | 2.00 | 2.42 |  | 296.00 | 0.48 | 0.93 | 8.26 | 4.83 | 1.67 |
| DM1-33 | 2.00 | 45.70 |  | 401.00 | 1.70 | 1.42 | 15.31 | 3.46 | 3.87 |
| DM1-34 | 4.00 | 24.55 |  | 352.00 | 1.63 | 1.52 | 10.54 | 3.92 | 2.50 |
| DM1-35 | 5.00 | 10.51 |  |  | 1.44 | 0.96 | 27.09 | 9.07 |  |
| DM1-36 | 3.00 | 13.29 |  |  | 1.44 | 1.24 | 33.63 | 5.23 | 2.15 |
| DM1-37 | 3.00 | 7.17 |  | 567.00 |  | 1.98 | 20.53 | 19.66 | 11.00 |
| DM1-38 |  |  |  |  |  |  |  |  |  |
| DM1-39 | 3.00 |  | 18.86 |  | 2.16 | 2.78 | 37.73 | 18.94 | 14.07 |
| DM1-40 | 1.00 | 2.59 | 20.24 |  | 1.30 | 1.29 | 14.71 | 15.29 | 1.98 |
| DM1-41 | 5.00 | 2.32 | 22.46 |  | 1.69 |  | 16.50 | 17.91 | 11.44 |
| DM1-42 | 1.00 | 2.01 | 15.90 |  | 2.21 | 3.45 | 26.58 | 24.96 | 15.92 |
| DM1-43 | 2.00 | 1.97 | 19.98 |  | 3.64 | 3.56 | 28.59 | 22.82 | 13.54 |
| DM1-44 | 6.00 | 3.92 | 26.90 |  | 1.78 |  | 21.60 | 4.72 | 5.06 |
| DM1-45 | 3.00 | 5.00 | 20.25 |  | 2.38 | 2.97 | 26.27 | 15.60 | 20.53 |
| DM1-46 | 2.00 | 1.37 | 20.53 |  | 2.86 | 3.76 | 39.99 | 42.99 | 18.13 |
| DM1-47 | 1.00 | 1.25 | 20.13 |  | 2.42 | 2.91 | 16.58 | 13.56 | 9.24 |
| DM1-48 | 2.00 | 1.03 | 19.63 |  | 2.72 | 3.60 | 26.12 | 23.87 | 10.52 |
| DM1-49 | 5.00 | 3.06 | 18.97 |  | 2.25 | 2.50 | 23.64 | 7.00 | 8.94 |
| DM1-50 | 1.00 | 1.45 | 0.43 |  |  |  | 9.49 | 4.02 | 1.95 |
| DM1-51 | 2.00 | 4.36 | 20.06 |  | 1.94 | 2.58 | 14.04 | 13.69 | 8.61 |
| DM1-52 | 6.00 | 3.82 | 18.57 |  | 1.80 | 2.25 | 18.39 | 9.85 | 5.68 |
| DM1-53 | 1.00 | 4.85 | 21.89 |  | 3.13 | 4.15 | 41.13 | 10.28 | 17.86 |
| DM1-54 | 3.00 | 4.24 | 18.00 |  | 1.58 | 3.04 | 18.30 | 23.68 | 15.42 |
| DM1-55 | 5.00 | 1.24 | 19.61 | 315.00 | 0.81 | 1.69 | 27.91 | 19.47 | 18.11 |
| DM1-56 | 4.00 | 8.11 | 22.63 | 360.00 | 1.68 | 2.46 | 19.50 | 15.56 | 7.16 |
| DM1-57 | 2.00 | 0.62 | 37.65 | 443.00 | 1.66 | 2.22 | 33.41 | 33.22 | 3.92 |
| DM1-58 | 5.00 |  | 18.16 |  | 1.96 | 2.94 | 38.28 | 18.73 | 14.08 |
| DM1-59 | 2.00 | 2.70 | 17.31 |  | 1.40 |  | 19.46 | 14.58 | 1.46 |
| DM1-60 | 5.00 |  | 20.35 |  | 1.67 |  | 21.95 | 19.43 | 11.56 |
| DM1-61 |  |  |  |  |  |  |  |  |  |
| DM1-62 | 5.00 | 10.03 | 27.10 |  | 1.80 |  | 26.57 | 3.27 | 2.96 |
| DM1-63 | 3.00 | 3.57 | 16.63 |  | 2.25 | 3.05 | 32.35 | 17.25 | 14.63 |
| DM1-64 | 1.00 |  | 21.30 |  | 2.61 | 3.57 | 42.03 | 46.01 | 19.31 |
| DM1-65 | 1.00 |  | 19.82 |  | 2.50 | 2.91 | 15.22 | 12.49 | 10.05 |
| DM1-66 | 1.00 |  | 19.07 |  | 2.33 | 3.60 | 24.01 | 20.74 | 14.12 |
| DM1-67 | 6.00 | 0.90 | 48.99 |  |  |  | 8.53 | 3.87 | 3.52 |
| DM1-68 | 5.00 | 2.91 | 17.96 | 488.00 | 2.25 | 2.73 | 11.51 | 9.52 | 3.26 |
| DM1-69 | 2.00 | 9.03 | 22.66 | 422.00 | 2.06 | 3.30 | 44.89 | 8.34 | 14.49 |
| DM1-70 | 5.00 | 1.95 | 19.49 | 562.00 | 1.88 | 3.05 | 21.36 | 23.53 | 11.55 |
| DM1-71 | 3.00 | 5.68 | 29.22 | 141.00 | 0.41 |  | 27.78 | 18.48 | 12.33 |
| DM1-72 | 3.00 | 9.75 | 20.97 | 417.00 | 1.80 | 2.56 | 20.49 | 14.23 | 8.96 |
| DM1-73 | 2.00 |  | 35.90 | 422.00 | 1.69 | 1.86 | 37.64 | 35.25 | 12.54 |
| DM1-74 | 2.00 | 2.48 | 19.37 |  | 1.80 | 2.67 | 29.42 | 20.34 | 10.86 |
| DM1-75 | 1.00 | 2.59 | 21.50 |  | 1.13 |  | 16.68 | 14.98 | 2.04 |
| DM1-76 | 5.00 | 10.13 | 22.52 |  | 1.71 | 2.22 | 19.89 | 19.44 | 11.41 |
| DM1-77 | 3.00 | 8.10 | 17.83 |  | 2.50 | 3.68 | 28.54 | 22.59 | 10.68 |
| DM1-78 | 4.00 | 4.47 | 16.69 |  | 1.91 | 2.83 | 27.39 | 15.21 | 15.29 |
| DM1-79 | 2.00 | 0.86 | 22.44 |  | 2.47 | 3.51 | 38.23 | 41.61 | 17.96 |
| DM1-80 | 1.00 | 1.04 | 18.39 |  | 2.61 | 2.99 | 15.57 | 12.87 | 12.15 |
| DM1-81 | 2.00 | 1.86 | 20.91 |  | 2.80 | 4.00 | 24.52 | 23.06 | 13.00 |
| DM1-82 | 1.00 | 0.77 | 0.43 |  |  |  | 10.53 | 4.46 | 4.15 |
| DM1-83 | 3.00 | 5.71 | 19.01 | 520.00 | 1.88 | 2.43 | 15.15 | 14.55 | 9.71 |
| DM1-84 | 3.00 | 3.89 | 18.16 | 454.00 | 1.37 | 2.67 | 17.86 | 11.11 | 5.37 |
| DM1-85 | 1.00 | 6.63 | 26.69 | 265.00 | 2.61 | 3.60 | 41.93 | 6.95 | 18.72 |
| DM1-86 | 4.00 |  | 17.29 | 553.00 | 1.90 | 3.10 | 23.60 | 25.05 | 12.70 |
| DM1-87 | 3.00 | 9.07 | 21.33 | 206.00 | 1.49 | 2.62 | 26.90 | 31.69 | 7.47 |
| DM1-88 | 2.00 | 1.57 | 33.61 | 392.00 | 1.73 | 1.90 |  | 37.55 | 11.95 |
