## Supplemental Table 3 for "RNA mis-splicing in children with myotonic dystrophy is associated with physical function"

| **Outcome variable** | **[MBNL]inferred – DM1 and CDM** |  |  |  |  |  |  |
| --- | --- | --- | --- | --- | --- | --- | --- |
| **Model 1** |  |  |  |  |  |  |  |
| **Analysis of Variance** | **SS** | **DF** | **MS** | **F (DFn, DFd)** | **P value** |  |  |
| Regression | 2.419 | 4 | 0.6047 | F (4, 40) = 23.57 | P<0.0001 |  |  |
| % Predicted 6MW | 0.0006540 | 1 | 0.0006540 | F (1, 40) = 0.02549 | P=0.8740 |  |  |
| % Predicted Walk/Run Velocity | 0.6538 | 1 | 0.6538 | F (1, 40) = 25.48 | P<0.0001 |  |  |
| % Predicted KE | 0.1072 | 1 | 0.1072 | F (1, 40) = 4.179 | P=0.0475 |  |  |
| % Predicted ADF | 0.1066 | 1 | 0.1066 | F (1, 40) = 4.154 | P=0.0482 |  |  |
| Residual | 1.026 | 40 | 0.02566 |  |  |  |  |
| Total | 3.445 | 44 |  |  |  |  |  |
| **Parameter estimates** | **Variable** | **Estimate** | **Standard error** | **95% CI (asymptotic)** | **\|t\|** | **P value** | **P value summary** |
| β0 | Intercept | -0.1162 | 0.1085 | -0.3355 to 0.1031 | 1.071 | 0.2908 | ns |
| β1 | % Predicted 6MW | 0.0003441 | 0.002155 | -0.004012 to 0.004700 | 0.1597 | 0.8740 | ns |
| β2 | % Predicted Walk/Run Velocity | 0.004033 | 0.0007990 | 0.002418 to 0.005648 | 5.048 | <0.0001 | **** |
| β3 | % Predicted KE | 0.0008911 | 0.0004359 | 1.014e-005 to 0.001772 | 2.044 | 0.0475 | * |
| β4 | % Predicted ADF | 0.002795 | 0.001371 | 2.345e-005 to 0.005567 | 2.038 | 0.0482 | * |
| **Goodness of Fit** |  |  |  |  |  |  |  |
| Degrees of Freedom | 40 |  |  |  |  |  |  |
| Multiple R | 0.8379 |  |  |  |  |  |  |
| R squared | 0.7021 |  |  |  |  |  |  |
| Adjusted R squared | 0.6723 |  |  |  |  |  |  |
| Sum of Squares | 1.026 |  |  |  |  |  |  |
| Sy.x | 0.1602 |  |  |  |  |  |  |
| RMSE | 0.1527 |  |  |  |  |  |  |
| AICc | -155.9 |  |  |  |  |  |  |
| **Multicollinearity** | **Variable** | **VIF** | **R2 with other variables** |  |  |  |  |
| β0 | Intercept |  |  |  |  |  |  |
| β1 | % Predicted 6MW | 1.672 | 0.4019 |  |  |  |  |
| β2 | % Predicted Walk/Run Velocity | 1.556 | 0.3575 |  |  |  |  |
| β3 | % Predicted KE | 1.586 | 0.3697 |  |  |  |  |
| β4 | % Predicted ADF | 1.825 | 0.4521 |  |  |  |  |
| **Normality of Residuals** | **Statistics** | **P value** | **Passed normality test (alpha=0.05)?** | **P value summary** |  |  |  |
| D'Agostino-Pearson omnibus (K2) | 3.609 | 0.1645 | Yes | ns |  |  |  |
| Anderson-Darling (A2*) | 0.3599 | 0.4339 | Yes | ns |  |  |  |
| Shapiro-Wilk (W) | 0.9727 | 0.3623 | Yes | ns |  |  |  |
| Kolmogorov-Smirnov (distance) | 0.07941 | >0.1000 | Yes | ns |  |  |  |
| **Data summary** |  |  |  |  |  |  |  |
| Rows analyzed (# cases) | 45 |  |  |  |  |  |  |
