## Supplemental Table 4 for "RNA mis-splicing in children with myotonic dystrophy is associated with physical function"

| **Outcome variable** | **[MBNL]inferred - CDM** |  |  |  |  |  |  |
| --- | --- | --- | --- | --- | --- | --- | --- |
| **Model 2** |  |  |  |  |  |  |  |
| **Analysis of Variance** | **SS** | **DF** | **MS** | **F (DFn, DFd)** | **P value** |  |  |
| Regression | 0.2828 | 3 | 0.09426 | F (3, 9) = 6.697 | P=0.0114 |  |  |
| % Predicted Walk/Run Velocity | 0.02891 | 1 | 0.02891 | F (1, 9) = 2.054 | P=0.1856 |  |  |
| % Predicted 9HPT | 0.008995 | 1 | 0.008995 | F (1, 9) = 0.6391 | P=0.4446 |  |  |
| % Predicted KE | 0.1076 | 1 | 0.1076 | F (1, 9) = 7.644 | P=0.0219 |  |  |
| Residual | 0.1267 | 9 | 0.01408 |  |  |  |  |
| Total | 0.4094 | 12 |  |  |  |  |  |
| **Parameter estimates** | **Variable** | **Estimate** | **Standard error** | **95% CI (asymptotic)** | **\|t\|** | **P value** | **P value summary** |
| β0 | Intercept | 0.3704 | 0.2742 | -0.2499 to 0.9908 | 1.351 | 0.2097 | ns |
| β1 | % Predicted Walk/Run Velocity | 0.001678 | 0.001171 | -0.0009708 to 0.004327 | 1.433 | 0.1856 | ns |
| β2 | % Predicted 9HPT | -0.0005758 | 0.0007203 | -0.002205 to 0.001054 | 0.7994 | 0.4446 | ns |
| β3 | % Predicted KE | 0.001659 | 0.0006000 | 0.0003016 to 0.003016 | 2.765 | 0.0219 | * |
| **Goodness of Fit** |  |  |  |  |  |  |  |
| Degrees of Freedom | 9 |  |  |  |  |  |  |
| Multiple R | 0.8310 |  |  |  |  |  |  |
| R squared | 0.6906 |  |  |  |  |  |  |
| Adjusted R squared | 0.5875 |  |  |  |  |  |  |
| Sum of Squares | 0.1267 |  |  |  |  |  |  |
| Sy.x | 0.1186 |  |  |  |  |  |  |
| RMSE | 0.1027 |  |  |  |  |  |  |
| AICc | -41.63 |  |  |  |  |  |  |
| **Multicollinearity** | **Variable** | **VIF** | **R2 with other variables** |  |  |  |  |
| β0 | Intercept |  |  |  |  |  |  |
| β1 | % Predicted Walk/Run Velocity | 1.801 | 0.4449 |  |  |  |  |
| β2 | % Predicted 9HPT | 1.980 | 0.4950 |  |  |  |  |
| β3 | % Predicted KE | 1.134 | 0.1184 |  |  |  |  |
| **Normality of Residuals** | **Statistics** | **P value** | **Passed normality test (alpha=0.05)?** | **P value summary** |  |  |  |
| D'Agostino-Pearson omnibus (K2) | 1.868 | 0.3931 | Yes | ns |  |  |  |
| Anderson-Darling (A2*) | 0.5226 | 0.1481 | Yes | ns |  |  |  |
| Shapiro-Wilk (W) | 0.9043 | 0.1532 | Yes | ns |  |  |  |
| Kolmogorov-Smirnov (distance) | 0.1968 | >0.1000 | Yes | ns |  |  |  |
| **Data summary** |  |  |  |  |  |  |  |
| Rows analyzed (# cases) | 13 |  |  |  |  |  |  |
