## Supplemental Table 5 for "RNA mis-splicing in children with myotonic dystrophy is associated with physical function"

| **Outcome variable** | **Walk/ Run Velocity (% Predicted) – Adults** |  |  |  |  |  |  |
| --- | --- | --- | --- | --- | --- | --- | --- |
| **Model 1** |  |  |  |  |  |  |  |
| **Analysis of Variance** | **SS** | **DF** | **MS** | **F (DFn, DFd)** | **P value** |  |  |
| Regression | 2882 | 2 | 1441 | F (2, 9) = 12.73 | P=0.0024 |  |  |
| Baseline [MBNL]inferred | 255.0 | 1 | 255.0 | F (1, 9) = 2.254 | P=0.1675 |  |  |
| Baseline % Predicted Walk/Run Velocity | 2844 | 1 | 2844 | F (1, 9) = 25.13 | P=0.0007 |  |  |
| Residual | 1018 | 9 | 113.2 |  |  |  |  |
| Total | 3901 | 11 |  |  |  |  |  |
| **Parameter estimates** | **Variable** | **Estimate** | **Standard error** | **95% CI (asymptotic)** | **\|t\|** | **P value** | **P value summary** |
| β0 | Intercept | 39.66 | 20.29 | -6.250 to 85.56 | 1.954 | 0.0824 | ns |
| β1 | Baseline [MBNL]inferred | -31.03 | 20.67 | -77.78 to 15.73 | 1.501 | 0.1675 | ns |
| β2 | Baseline % Predicted Walk/Run Velocity | 0.8320 | 0.1660 | 0.4566 to 1.207 | 5.013 | 0.0007 | *** |
| **Goodness of Fit** |  |  |  |  |  |  |  |
| Degrees of Freedom | 9 |  |  |  |  |  |  |
| Multiple R | 0.8596 |  |  |  |  |  |  |
| R squared | 0.7389 |  |  |  |  |  |  |
| Adjusted R squared | 0.6809 |  |  |  |  |  |  |
| Sum of Squares | 1018 |  |  |  |  |  |  |
| Sy.x | 10.64 |  |  |  |  |  |  |
| RMSE | 9.622 |  |  |  |  |  |  |
| AICc | 67.01 |  |  |  |  |  |  |
| **Multicollinearity** | **Variable** | **VIF** | **R2 with other variables** |  |  |  |  |
| β0 | Intercept |  |  |  |  |  |  |
| β1 | Baseline [MBNL]inferred | 1.196 | 0.1642 |  |  |  |  |
| β2 | Baseline % Predicted Walk/Run Velocity | 1.196 | 0.1642 |  |  |  |  |
| **Normality of Residuals** | **Statistics** | **P value** | **Passed normality test (alpha=0.05)?** | **P value summary** |  |  |  |
| D'Agostino-Pearson omnibus (K2) | 0.9126 | 0.6336 | Yes | ns |  |  |  |
| Anderson-Darling (A2*) | 0.4299 | 0.2562 | Yes | ns |  |  |  |
| Shapiro-Wilk (W) | 0.9289 | 0.3688 | Yes | ns |  |  |  |
| Kolmogorov-Smirnov (distance) | 0.2152 | >0.1000 | Yes | ns |  |  |  |
| **Data summary** |  |  |  |  |  |  |  |
| Rows analyzed (# cases) | 12 |  |  |  |  |  |  |
| **Outcome variable** | **Walk/ Run Velocity (% Predicted) – Children** |  |  |  |  |  |  |
| **Model 2** |  |  |  |  |  |  |  |
| **Analysis of Variance** | **SS** | **DF** | **MS** | **F (DFn, DFd)** | **P value** |  |  |
| Regression | 12203 | 2 | 6101 | F (2, 11) = 12.84 | P=0.0013 |  |  |
| Baseline [MBNL]inferred | 67.95 | 1 | 67.95 | F (1, 11) = 0.1430 | P=0.7125 |  |  |
| Baseline % Predicted Walk/Run Velocity | 9156 | 1 | 9156 | F (1, 11) = 19.27 | P=0.0011 |  |  |
| Residual | 5226 | 11 | 475.1 |  |  |  |  |
| Total | 17428 | 13 |  |  |  |  |  |
| **Parameter estimates** | **Variable** | **Estimate** | **Standard error** | **95% CI (asymptotic)** | **\|t\|** | **P value** | **P value summary** |
| β0 | Intercept | -8.508 | 25.66 | -64.98 to 47.97 | 0.3316 | 0.7464 | ns |
| β1 | Baseline [MBNL]inferred | 13.24 | 35.00 | -63.80 to 90.27 | 0.3782 | 0.7125 | ns |
| β2 | Baseline % Predicted Walk/Run Velocity | 1.035 | 0.2357 | 0.5160 to 1.553 | 4.390 | 0.0011 | ** |
| **Goodness of Fit** |  |  |  |  |  |  |  |
| Degrees of Freedom | 11 |  |  |  |  |  |  |
| Multiple R | 0.8368 |  |  |  |  |  |  |
| R squared | 0.7002 |  |  |  |  |  |  |
| Adjusted R squared | 0.6456 |  |  |  |  |  |  |
| Sum of Squares | 5226 |  |  |  |  |  |  |
| Sy.x | 21.80 |  |  |  |  |  |  |
| RMSE | 20.05 |  |  |  |  |  |  |
| AICc | 95.36 |  |  |  |  |  |  |
| **Multicollinearity** | **Variable** | **VIF** | **R2 with other variables** |  |  |  |  |
| β0 | Intercept |  |  |  |  |  |  |
| β1 | Baseline [MBNL]inferred | 1.232 | 0.1880 |  |  |  |  |
| β2 | Baseline % Predicted Walk/Run Velocity | 1.232 | 0.1880 |  |  |  |  |
| **Normality of Residuals** | **Statistics** | **P value** | **Passed normality test (alpha=0.05)?** | **P value summary** |  |  |  |
| D'Agostino-Pearson omnibus (K2) | 10.76 | 0.0046 | No | ** |  |  |  |
| Anderson-Darling (A2*) | 1.004 | 0.0083 | No | ** |  |  |  |
| Shapiro-Wilk (W) | 0.8364 | 0.0146 | No | * |  |  |  |
| Kolmogorov-Smirnov (distance) | 0.2504 | 0.0175 | No | * |  |  |  |
| **Data summary** |  |  |  |  |  |  |  |
| Rows analyzed (# cases) | 14 |  |  |  |  |  |  |
