## Supplemental Table 6 for "RNA mis-splicing in children with myotonic dystrophy is associated with physical function"

| **Outcome variable** | **12-month Ankle Dorsiflexion Strength (% Predicted) - Adults** |  |  |  |  |  |  |
| --- | --- | --- | --- | --- | --- | --- | --- |
| **Model 1** |  |  |  |  |  |  |  |
| **Analysis of Variance** | **SS** | **DF** | **MS** | **F (DFn, DFd)** | **P value** |  |  |
| Regression | 4931 | 2 | 2466 | F (2, 12) = 18.44 | P=0.0002 |  |  |
| Baseline [MBNL]inferred | 456.4 | 1 | 456.4 | F (1, 12) = 3.413 | P=0.0895 |  |  |
| Baseline % Predicted Ankle Dorsiflexion | 3021 | 1 | 3021 | F (1, 12) = 22.59 | P=0.0005 |  |  |
| Residual | 1605 | 12 | 133.7 |  |  |  |  |
| Total | 6536 | 14 |  |  |  |  |  |
| **Parameter estimates** | **Variable** | **Estimate** | **Standard error** | **95% CI (asymptotic)** | **\|t\|** | **P value** | **P value summary** |
| β0 | Intercept | 8.159 | 12.25 | -18.54 to 34.86 | 0.6658 | 0.5181 | ns |
| β1 | Baseline [MBNL]inferred | 38.58 | 20.89 | -6.921 to 84.09 | 1.847 | 0.0895 | ns |
| β2 | Baseline % Predicted Ankle Dorsiflexion | 0.4888 | 0.1028 | 0.2647 to 0.7128 | 4.753 | 0.0005 | *** |
| **Goodness of Fit** |  |  |  |  |  |  |  |
| Degrees of Freedom | 12 |  |  |  |  |  |  |
| Multiple R | 0.8686 |  |  |  |  |  |  |
| R squared | 0.7545 |  |  |  |  |  |  |
| Adjusted R squared | 0.7136 |  |  |  |  |  |  |
| Sum of Squares | 1605 |  |  |  |  |  |  |
| Sy.x | 11.56 |  |  |  |  |  |  |
| RMSE | 10.71 |  |  |  |  |  |  |
| AICc | 82.09 |  |  |  |  |  |  |
| **Multicollinearity** | **Variable** | **VIF** | **R2 with other variables** |  |  |  |  |
| β0 | Intercept |  |  |  |  |  |  |
| β1 | Baseline [MBNL]inferred | 1.144 | 0.1259 |  |  |  |  |
| β2 | Baseline % Predicted Ankle Dorsiflexion | 1.144 | 0.1259 |  |  |  |  |
| **Normality of Residuals** | **Statistics** | **P value** | **Passed normality test (alpha=0.05)?** | **P value summary** |  |  |  |
| D'Agostino-Pearson omnibus (K2) | 0.4655 | 0.7923 | Yes | ns |  |  |  |
| Anderson-Darling (A2*) | 0.2675 | 0.6331 | Yes | ns |  |  |  |
| Shapiro-Wilk (W) | 0.9599 | 0.6905 | Yes | ns |  |  |  |
| Kolmogorov-Smirnov (distance) | 0.1441 | >0.1000 | Yes | ns |  |  |  |
| **Data summary** |  |  |  |  |  |  |  |
| Rows analyzed (# cases) | 15 |  |  |  |  |  |  |
| **Outcome variable** | **12-month Ankle Dorsiflexion Strength (% Predicted) - Children** |  |  |  |  |  |  |
| **Model 2** |  |  |  |  |  |  |  |
| **Analysis of Variance** | **SS** | **DF** | **MS** | **F (DFn, DFd)** | **P value** |  |  |
| Regression | 883.6 | 2 | 441.8 | F (2, 2) = 0.3419 | P=0.7452 |  |  |
| Baseline [MBNL]inferred | 15.31 | 1 | 15.31 | F (1, 2) = 0.01185 | P=0.9233 |  |  |
| Baseline % Predicted Ankle Dorsiflexion | 810.7 | 1 | 810.7 | F (1, 2) = 0.6273 | P=0.5114 |  |  |
| Residual | 2584 | 2 | 1292 |  |  |  |  |
| Total | 3468 | 4 |  |  |  |  |  |
| **Parameter estimates** | **Variable** | **Estimate** | **Standard error** | **95% CI (asymptotic)** | **\|t\|** | **P value** | **P value summary** |
| β0 | Intercept | 16.73 | 94.52 | -390.0 to 423.4 | 0.1770 | 0.8758 | ns |
| β1 | Baseline [MBNL]inferred | 12.40 | 113.9 | -477.9 to 502.7 | 0.1089 | 0.9233 | ns |
| β2 | Baseline % Predicted Ankle Dorsiflexion | 0.6690 | 0.8446 | -2.965 to 4.303 | 0.7920 | 0.5114 | ns |
| **Goodness of Fit** |  |  |  |  |  |  |  |
| Degrees of Freedom | 2 |  |  |  |  |  |  |
| Multiple R | 0.5048 |  |  |  |  |  |  |
| R squared | 0.2548 |  |  |  |  |  |  |
| Adjusted R squared | -0.4904 |  |  |  |  |  |  |
| Sum of Squares | 2584 |  |  |  |  |  |  |
| Sy.x | 35.95 |  |  |  |  |  |  |
| RMSE | 25.42 |  |  |  |  |  |  |
| AICc |  |  |  |  |  |  |  |
| **Multicollinearity** | **Variable** | **VIF** | **R2 with other variables** |  |  |  |  |
| β0 | Intercept |  |  |  |  |  |  |
| β1 | Baseline [MBNL]inferred | 1.026 | 0.02519 |  |  |  |  |
| β2 | Baseline % Predicted Ankle Dorsiflexion | 1.026 | 0.02519 |  |  |  |  |
| **Normality of Residuals** | **Statistics** | **P value** | **Passed normality test (alpha=0.05)?** | **P value summary** |  |  |  |
| D'Agostino-Pearson omnibus (K2) | N too small |  |  |  |  |  |  |
| Anderson-Darling (A2*) | N too small |  |  |  |  |  |  |
| Shapiro-Wilk (W) | 0.9893 | 0.9770 | Yes | ns |  |  |  |
| Kolmogorov-Smirnov (distance) | 0.1696 | >0.1000 | Yes | ns |  |  |  |
| **Data summary** |  |  |  |  |  |  |  |
| Rows analyzed (# cases) | 5 |  |  |  |  |  |  |
