## Supplemental Table 7 for "RNA mis-splicing in children with myotonic dystrophy is associated with physical function"

| **Outcome variable** | **12-month Stair Climbing Velocity (% Predicted) - Adults and Children** |  |  |  |  |  |  |
| --- | --- | --- | --- | --- | --- | --- | --- |
| **Model 1** |  |  |  |  |  |  |  |
| **Analysis of Variance** | **SS** | **DF** | **MS** | **F (DFn, DFd)** | **P value** |  |  |
| Regression | 7207 | 2 | 3604 | F (2, 25) = 56.98 | P<0.0001 |  |  |
| Baseline [MBNL]inferred | 440.3 | 1 | 440.3 | F (1, 25) = 6.961 | P=0.0141 |  |  |
| Baseline % Predicted Stair Velocity | 6423 | 1 | 6423 | F (1, 25) = 101.6 | P<0.0001 |  |  |
| Residual | 1581 | 25 | 63.25 |  |  |  |  |
| Total | 8789 | 27 |  |  |  |  |  |
| **Parameter estimates** | **Variable** | **Estimate** | **Standard error** | **95% CI (asymptotic)** | **\|t\|** | **P value** | **P value summary** |
| β0 | Intercept | -0.5239 | 6.848 | -14.63 to 13.58 | 0.07651 | 0.9396 | ns |
| β1 | Baseline [MBNL]inferred | 23.11 | 8.759 | 5.070 to 41.15 | 2.638 | 0.0141 | * |
| β2 | Baseline % Predicted Stair Velocity | 0.7103 | 0.07049 | 0.5652 to 0.8555 | 10.08 | <0.0001 | **** |
| **Goodness of Fit** |  |  |  |  |  |  |  |
| Degrees of Freedom | 25 |  |  |  |  |  |  |
| Multiple R | 0.9056 |  |  |  |  |  |  |
| R squared | 0.8201 |  |  |  |  |  |  |
| Adjusted R squared | 0.8057 |  |  |  |  |  |  |
| Sum of Squares | 1581 |  |  |  |  |  |  |
| Sy.x | 7.953 |  |  |  |  |  |  |
| RMSE | 7.653 |  |  |  |  |  |  |
| AICc | 122.7 |  |  |  |  |  |  |
| **Multicollinearity** | **Variable** | **VIF** | **R2 with other variables** |  |  |  |  |
| β0 | Intercept |  |  |  |  |  |  |
| β1 | Baseline [MBNL]inferred | 1.008 | 0.007451 |  |  |  |  |
| β2 | Baseline % Predicted Stair Velocity | 1.008 | 0.007451 |  |  |  |  |
| **Normality of Residuals** | **Statistics** | **P value** | **Passed normality test (alpha=0.05)?** | **P value summary** |  |  |  |
| D'Agostino-Pearson omnibus (K2) | 1.533 | 0.4646 | Yes | ns |  |  |  |
| Anderson-Darling (A2*) | 0.2042 | 0.8604 | Yes | ns |  |  |  |
| Shapiro-Wilk (W) | 0.9788 | 0.8215 | Yes | ns |  |  |  |
| Kolmogorov-Smirnov (distance) | 0.09382 | >0.1000 | Yes | ns |  |  |  |
| **Data summary** |  |  |  |  |  |  |  |
| Rows analyzed (# cases) | 28 |  |  |  |  |  |  |
| **Outcome variable** | **12-month Stair Climbing Velocity (% Predicted) - Children** |  |  |  |  |  |  |
| **Model 2** |  |  |  |  |  |  |  |
| **Analysis of Variance** | **SS** | **DF** | **MS** | **F (DFn, DFd)** | **P value** |  |  |
| Regression | 3695 | 2 | 1847 | F (2, 11) = 41.90 | P<0.0001 |  |  |
| Baseline [MBNL]inferred | 137.7 | 1 | 137.7 | F (1, 11) = 3.123 | P=0.1049 |  |  |
| Baseline % Predicted Stair Velocity | 3198 | 1 | 3198 | F (1, 11) = 72.53 | P<0.0001 |  |  |
| Residual | 485.0 | 11 | 44.09 |  |  |  |  |
| Total | 4180 | 13 |  |  |  |  |  |
| **Parameter estimates** | **Variable** | **Estimate** | **Standard error** | **95% CI (asymptotic)** | **\|t\|** | **P value** | **P value summary** |
| β0 | Intercept | -6.398 | 7.724 | -23.40 to 10.60 | 0.8284 | 0.4251 | ns |
| β1 | Baseline [MBNL]inferred | 17.26 | 9.768 | -4.238 to 38.76 | 1.767 | 0.1049 | ns |
| β2 | Baseline % Predicted Stair Velocity | 0.9154 | 0.1075 | 0.6788 to 1.152 | 8.516 | <0.0001 | **** |
| **Goodness of Fit** |  |  |  |  |  |  |  |
| Degrees of Freedom | 11 |  |  |  |  |  |  |
| Multiple R | 0.9402 |  |  |  |  |  |  |
| R squared | 0.8840 |  |  |  |  |  |  |
| Adjusted R squared | 0.8629 |  |  |  |  |  |  |
| Sum of Squares | 485.0 |  |  |  |  |  |  |
| Sy.x | 6.640 |  |  |  |  |  |  |
| RMSE | 6.108 |  |  |  |  |  |  |
| AICc | 62.08 |  |  |  |  |  |  |
| **Multicollinearity** | **Variable** | **VIF** | **R2 with other variables** |  |  |  |  |
| β0 | Intercept |  |  |  |  |  |  |
| β1 | Baseline [MBNL]inferred | 1.034 | 0.03249 |  |  |  |  |
| β2 | Baseline % Predicted Stair Velocity | 1.034 | 0.03249 |  |  |  |  |
| **Normality of Residuals** | **Statistics** | **P value** | **Passed normality test (alpha=0.05)?** | **P value summary** |  |  |  |
| D'Agostino-Pearson omnibus (K2) | 0.06238 | 0.9693 | Yes | ns |  |  |  |
| Anderson-Darling (A2*) | 0.1358 | 0.9695 | Yes | ns |  |  |  |
| Shapiro-Wilk (W) | 0.9901 | 0.9996 | Yes | ns |  |  |  |
| Kolmogorov-Smirnov (distance) | 0.1066 | >0.1000 | Yes | ns |  |  |  |
| **Data summary** |  |  |  |  |  |  |  |
| Rows analyzed (# cases) | 14 |  |  |  |  |  |  |
| **Outcome variable** | **12-month Stair Climbing Speed (% Predicted) - Adults** |  |  |  |  |  |  |
| **Model 3** |  |  |  |  |  |  |  |
| **Analysis of Variance** | **SS** | **DF** | **MS** | **F (DFn, DFd)** | **P value** |  |  |
| Regression | 2264 | 2 | 1132 | F (2, 11) = 15.46 | P=0.0006 |  |  |
| Baseline [MBNL]inferred | 304.8 | 1 | 304.8 | F (1, 11) = 4.162 | P=0.0661 |  |  |
| Baseline % Predicted Stair Velocity | 1543 | 1 | 1543 | F (1, 11) = 21.07 | P=0.0008 |  |  |
| Residual | 805.6 | 11 | 73.23 |  |  |  |  |
| Total | 3070 | 13 |  |  |  |  |  |
| **Parameter estimates** | **Variable** | **Estimate** | **Standard error** | **95% CI (asymptotic)** | **\|t\|** | **P value** | **P value summary** |
| β0 | Intercept | 5.637 | 11.35 | -19.35 to 30.62 | 0.4966 | 0.6293 | ns |
| β1 | Baseline [MBNL]inferred | 30.98 | 15.19 | -2.443 to 64.41 | 2.040 | 0.0661 | ns |
| β2 | Baseline % Predicted Stair Velocity | 0.5539 | 0.1207 | 0.2883 to 0.8195 | 4.590 | 0.0008 | *** |
| **Goodness of Fit** |  |  |  |  |  |  |  |
| Degrees of Freedom | 11 |  |  |  |  |  |  |
| Multiple R | 0.8588 |  |  |  |  |  |  |
| R squared | 0.7376 |  |  |  |  |  |  |
| Adjusted R squared | 0.6899 |  |  |  |  |  |  |
| Sum of Squares | 805.6 |  |  |  |  |  |  |
| Sy.x | 8.558 |  |  |  |  |  |  |
| RMSE | 7.872 |  |  |  |  |  |  |
| AICc | 69.18 |  |  |  |  |  |  |
| **Multicollinearity** | **Variable** | **VIF** | **R2 with other variables** |  |  |  |  |
| β0 | Intercept |  |  |  |  |  |  |
| β1 | Baseline [MBNL]inferred | 1.052 | 0.04937 |  |  |  |  |
| β2 | Baseline % Predicted Stair Velocity | 1.052 | 0.04937 |  |  |  |  |
| **Normality of Residuals** | **Statistics** | **P value** | **Passed normality test (alpha=0.05)?** | **P value summary** |  |  |  |
| D'Agostino-Pearson omnibus (K2) | 2.508 | 0.2853 | Yes | ns |  |  |  |
| Anderson-Darling (A2*) | 0.6009 | 0.0952 | Yes | ns |  |  |  |
| Shapiro-Wilk (W) | 0.9054 | 0.1352 | Yes | ns |  |  |  |
| Kolmogorov-Smirnov (distance) | 0.1976 | >0.1000 | Yes | ns |  |  |  |
| **Data summary** |  |  |  |  |  |  |  |
| Rows analyzed (# cases) | 14 |  |  |  |  |  |  |
