## Supplementary material for "RNA mis-splicing in children with myotonic dystrophy is associated with physical function": DMCRN Consortium Members

| **DMCRN Consortium** | |
| --- | --- |
| **Affiliation(s)** | **Role: Name, Titles** |
| University of Rochester Medical Center | **PI:** Johanna Hamel, MD  Associate Professor  Department of Neurology  Neuromuscular Disease and Department of Pathology and Laboratory Medicine  **Primary CRC:** Jeanne Dekdebrun, MS  Senior Research Coordinator  **Primary CE:** Katy Eichinger, PT, PhD, DPT, NCS  Associate Professor  Department of Neurology  Neuromuscular Division |
| The Ohio State University Wexner Medical Center  Department of Neurology, Division of Neuromuscular Diseases | **PI:** Bakri Elsheikh, MBBS, FRCP, FAAN  Professor of Neurology  Director of the Neuromuscular Division  Director of OSU Global Neurology Initiative  Director of the OSU Wexner Medical Center Muscular Dystrophy Association Care Center  Director of Clinical Neurophysiology Fellowship  Department of Neurology, Neuromuscular Division  **CRC:** Kaneshia Hives  **CE:** Kathryn Jira, PT, DPT |
| University of Kansas Medical Center | **PI:** Jeffrey Statland, MD  **Primary CRC:** Kaylene Whited, MS  **Primary CE:** Sandhya Sasidharan, PT |
| University of Florida College of Medicine | **PI:** Sub Subramony, MD  Professor of Neurology and Pediatrics  University of Florida College of Medicine  **Primary CRC:** Alex Rodriguez  **Primary CE**: Donovan Lott, PT, PhD, CSCS  Research Professor  Department of Physical Therapy |
| University of California, Los Angeles | **PI:** Perry Shieh, MD, PhD  **Primary CRC:** Dennis Fernando  **Primary CE:** Christy Skura, PT, DPT, PCS |
| Stanford University  Department of Neurology & Neurological Services | **PI:** Jacinda Sampson, MD, PhD  **Primary CRC:** Sara Ismail  **Primary CE:** Tina Duong, PT, PhD |
| Houston Methodist Neurological Institute | **PI:** Ericka P. Greene, MD  **Primary CRC**: Aramide Balogun, BS  **Primary CE**: Dora Maldonado |
| University of Iowa | **PI:** Andrea Swenson, MD  **Primary CRC:** Maegan Tyrell, BA, CHES  **Primary CE:** Amy Yotty, PT, DPT  University of Iowa Health Care |
| Virginia Commonwealth University | **PI:** Nicholas Johnson, MD, M. Sci, FAAN  **Primary CRC:** Carino Jennings  **Primary CE:** Aileen Jones, PT, DPT |
| Radboud University Medical Center | **PI:** Karlien Mul, MD, PhD  **Primary CRC:** Monique Plieger  **Primary CE:** Judith van Engelen-Kanters |
| Fondazione Serena ONLUS - Centro Clinico NeMO Milano | **PI:** Valeria Sansone, MD, PhD  **Primary CRC:** Michela Nani, RN  **Primary CE:** Enrico Cossu, PT  Valentina Franchino, TNPEE |
| Friedrich Baur Institute, Department of Neurology, LMU University Hospital, LMU Munich | **PI:** Benedikt Schoser, FEAN  **Primary CRC/CE:** Corinna Wirner-Piotrowski, M Sc. |
| The National Hospital for Neurology and Neurosurgery, University College London Hospitals NHS Foundation Trust | **PI:** Chris Turner, MD, PhD, FRCP  Divisional Clinical Director, Queen Square.  Consultant Neurologist and UCL Honorary Clinical Associate Professor  University College Hospitals NHS Foundation Trust  **Primary CRC:** Nikoletta Nikolenko, MD, PhD  **Primary CE:** Charlotte Massey, BSc  Sheffield Institute for Translational Neuroscience (SITraN)  Department of Neuroscience, University of Sheffield |
| University of Colorado | **PI:** Thomas Ragole, MD  Assistant Professor  **Primary CRC**: Alyssa Avilez, BS  **Primary CE:** Talia Strahler, PT, DPT, OCS  Instructor |
| The University of Auckland, Centre for Brain Research Neurogenetics Clinic | **PI:** Richard Roxburgh, MBChB PhD FRACP  **Primary CRC**: Sarah Nagar, PGDipPH, BSc  **Primary CE:** Sarah Mollet, BHSc Physiotherapy  University of Auckland  CBR Neurogenetics Clinic |
| UC San Diego Health  Rady’s Children’s Hospital San Diego | **PI:** Chamindra Laverty, MD  **Primary CRC**: Mariah Stechschulte, BS  **Primary CE:** Kristine Negrete, DPT  Neurolab 360 |
| Atkinson-Morley Neuromuscular Centre  St George's University Hospitals NHS Foundation Trust and Neuroscience and Cell Biology Research Institute  St George's University of London | **PI:** Emma Matthews, MBChB, PhD, FRCP  **Primary CRC:** Claire Gilmartin  **Primary CE:** Claire O’Farrell, BSc |
| Groupe de recherche interdisciplinaire sur les maladies neuromusculaires (GRIMN)  Integrated University Health and Social Services Centres (CIUSSS) of Saguenay Lac-St-Jean  Sherbrooke University | **PI:** Cynthia Gagnon, OT, PhD  **Primary CRC:** Justine Dolbec, M. Sc.  **Primary CE:** Amelie Lemire, B. Sc. Kin. |
| Osaka University | **PI:** Masanori Takahashi, MD, PhD  Osaka University  **Primary CRC:** Shizuko Taima, RN  Aomori National Hospital  **Primary CE:** Suzuki Manabu, RPT  Aomori National Hospital |
| University of Texas Health, Antonio | **PI:** Matthew Wicklund  **Primary CRC:** TBD  **Primary CE:** TBD |
