## Supplementary figures and images for "RNA mis-splicing in children with myotonic dystrophy is associated with physical function"

### Supplemental Figure 1

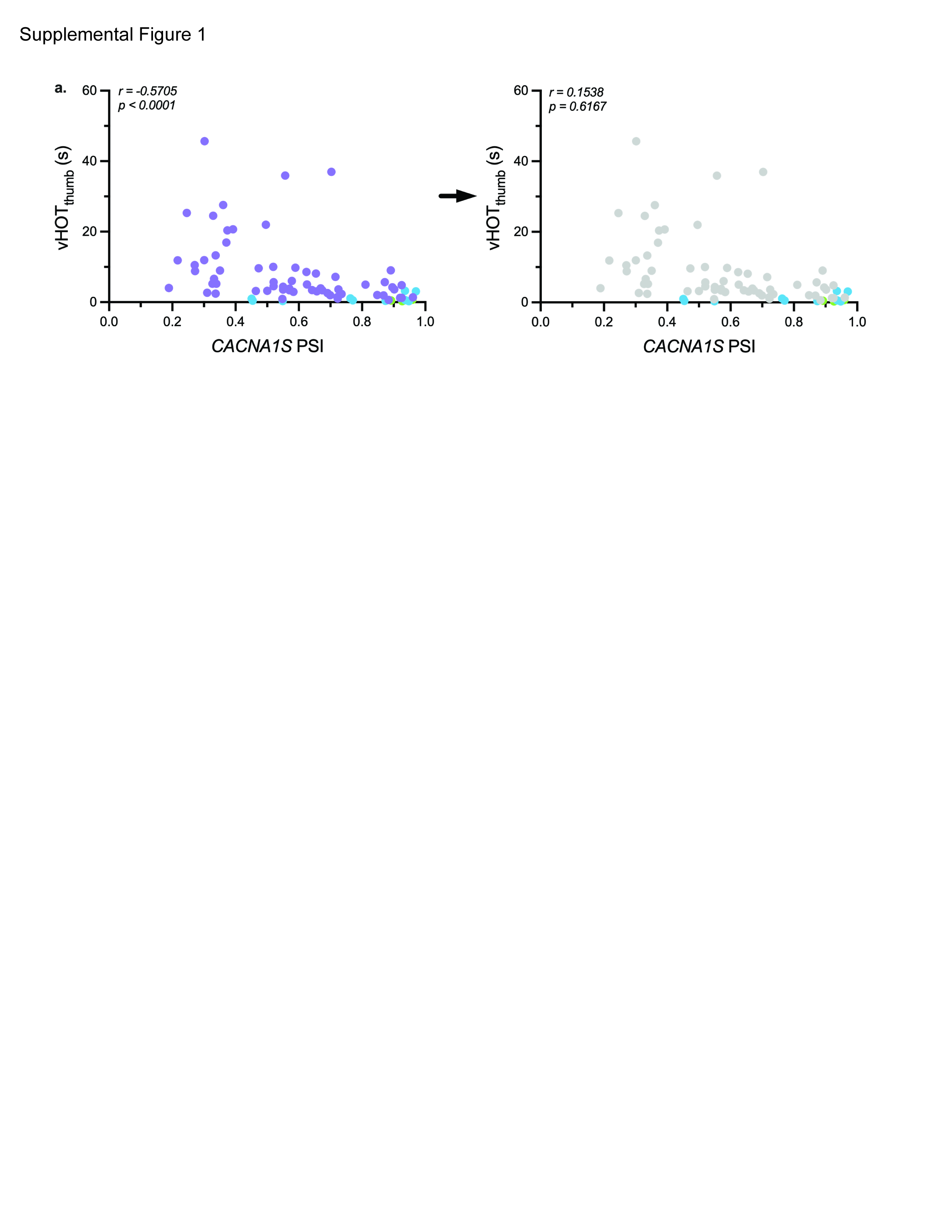
